# Insights into plastic biodegradation: community composition and functional capabilities of the superworm (*Zophobas morio*) microbiome in styrofoam feeding trials

**DOI:** 10.1101/2022.05.16.492041

**Authors:** Jiarui Sun, Apoorva Prabhu, Samuel Aroney, Christian Rinke

## Abstract

Plastics are inexpensive and widely used organic polymers, but their high durability hinders biodegradation. Polystyrene, including extruded polystyrene also known as styrofoam, is among the most commonly produced plastics worldwide and is recalcitrant to microbial degradation. In this study, we assessed changes in the gut microbiome of superworms (*Zophobas morio*) reared on bran, polystyrene, or under starvation conditions over a three weeks’ time period. Superworms on all diets were able to complete their life cycle to pupae and imago, although superworms reared on polystyrene had minimal weight gains, resulting in lower pupation rates. The change in microbial gut communities from baseline differed considerably between diet groups, with polystyrene and starvation groups characterized by a loss of microbial diversity and the presence of opportunistic pathogens. Inferred microbial functions enriched in the polystyrene group included transposon movements, membrane restructuring, and adaptations to oxidative stress. We detected several encoded enzymes with reported polystyrene and styrene degradation abilities, supporting previous reports of polystyrene degrading bacteria in the superworm gut. By recovering metagenome-assembled genomes (MAGs) we linked phylogeny and functions and identified genera including *Pseudomonas, Rhodococcus* and *Corynebacterium*, that possess genes associated with polystyrene degradation. In conclusion, our results provide the first metagenomic insights into the metabolic pathways used by the gut microbiome of superworms to degrade polystyrene. Our results also confirm that superworms can survive on polystyrene feed, however, this diet has considerable negative impacts on host gut microbiome diversity and health.

**Impact Statement:** Increasing plastic pollution is a major environmental problem, and a recently proposed way to counteract this trend is to embrace a circular economy, in which used materials are recycled, rather than disposed of. An important step to facilitate this process is to invent new approaches for upcycling of plastic waste to desirable consumer products. Microbial plastic degradation and conversion is likely to play a considerable part in shaping a circular economy, by engineering microbes or their enzymes to bio-upcycle plastic waste. A first step towards actualizing this goal is to identify microbes that can degrade polystyrene and to investigate the enzymes and pathways involved. Our study represents the first metagenomic analysis of an insect gut microbiome on a polystyrene diet. It identifies bacteria with polystyrene and styrene degrading abilities, and infers enzymes and pathways involved in these reactions. Therefore, our results contribute towards understanding microbial polystyrene degradation and will provide a base for future investigations into microbial upcycling of plastic waste.

## INTRODUCTION

Plastics are integral to the global economy and have become a part of our daily lives by providing excessive amounts of short-lived and disposable items (Thompson et al., 2009). Global plastic production reached nearly 360 million tonnes in 2018 and plastic demand is predicted to grow substantially over the next decade, while recycling rates are likely to remain low (Plastics Europe, 2019). This lack of recycling combined with the high durability of plastics has resulted in a wide range of negative environmental impacts (Barnes et al., 2009). One of the most prevalent polymers globally is the durable thermoplastic polystyrene (PS), which accounts for up to 7-10% of the total non-fiber plastic production (Geyer et al., 2017; Plastics Europe, 2016). Among the various PS types, PS foam is most widely used in consumer products (Ho et al., 2018) and is commonly referred to as styrofoam. Like other plastics, PS persists in nature for decades (Kaplan et al., 1979; Otake et al., 1995), and can potentially harm wildlife and subsequently humans (Barnes et al., 2009; Rochman et al., 2015). Historically PS was considered to be recalcitrant to microbes (Albertsson and Karlsson, 1990), and studies of PS degradation focused mainly on chemical and mechanistic forces (O’Brine and Thompson, 2010; Singh and Sharma, 2008). Indeed, the high molecular weight and hydrophobic character of PS polymers make them a difficult target for biodegradation since it prevents hydrolysis, which is the main pathway of natural polymer degradation (Krueger et al., 2017). However, initial abiotic degradation of PS has been reported to increase the polymer’s bioavailability in the environment by creating more exposed surfaces for biodegradation (Gewert et al., 2015; Singh and Sharma, 2008; Wei and Zimmermann, 2017). Over the last decades, several microbial isolates have been reported to utilize PS as a carbon source, albeit at a slow pace, with a reported PS weight loss ranging from 0.13 to 7% per day (**Table S1**). Thereby, environmental PS biodegradation is likely enhanced by cooperative processes involving microbial consortia. Indeed, mixed communities from soils and liquid enrichment cultures have been reported to degrade PS (Sielicki et al., 1978), and marine microbial consortia effectively reduced the weight of PS films suggesting that they are capable of degrading weathered PS pieces (Syranidou et al., 2017).

Insect gut microbiomes hold promising potential for efficient plastic degradation by combining both physical and biochemical mechanisms. The insect uses mechanical force, i.e. its oral appendages, to physically break up and ingest plastic particles which can subsequently be further degraded by the microbial community in the host’s digestive tract. Indeed, several recent studies have reported that insect microbiomes are capable of plastic degradation. Two bacterial strains, *Enterobacter asburiae* YT1 and *Bacillus sp*. YP1, isolated from the gut of waxworms, a common name for the larvae of the Indian mealmoth *Plodia interpunctella*, were capable of degrading polyethylene (PE) (Yang et al., 2014). Larvae of the wax moth, *Galleria mellonella*, have also been reported to digest PE, yet the role of its intestinal microbiome is not well understood (Bombelli et al., 2017), and the reported evidence for PE biodegradation, in particular the detection of ethylene glycol, has been called into question (Weber et al., 2017).

Currently, the best characterized plastic degrading insect gut microbiome is that of mealworms, a common name for the larvae of the mealworm beetle *Tenebrio molitor* (Tenebrionidae) (Y. Yang et al., 2015a, 2015b). Mealworms were able to survive on PS as a sole diet, and degraded long-chain PS molecules in their gut, resulting in the formation of depolymerized metabolites (Y. Yang et al., 2015b). The majority of PS was converted into CO2 (47.7%) and egested as feces (49.2%), with only a limited amount (0.5%) being incorporated into biomass. The gut microbiome was deemed essential for the reported PS breakdown since mealworms, treated with antibiotics, lost the ability to depolymerize PS (Y. Yang et al., 2015a). Furthermore, the Firmicute *Exiguobacterium sp*. strain YT2 isolated from a mealworm gut successfully formed a biofilm on PS and was able to degrade 7.4 ± 0.4% of the PS pieces over a 60-day incubation period (Y. Yang et al., 2015a). Another study, investigating the degradation of PE and mixed plastics in *T. molitor*, reported comparable biodegradation rates for PS and recovered two OTUs (*Citrobacter* sp. and *Kosakonia* sp.) associated with both PE and PS degradation (Brandon et al., 2018). The authors concluded that, based on the observed degradation of mixed polymers, plastic degradation within the mealworm gut is nonspecific in terms of plastic type.

Superworms, a common name for the larval stages of the darkling beetle *Zophobas morio* Fabricius, 1776 (Coleoptera, Tenebrionidae), have also been reported to survive on PS as the sole diet (Miao and Zhang, 2010; Yang et al., 2020). Recently, the larvae were found to convert 36.7% of the ingested PS carbon to CO_2_ in a study using the heterotypic synonym *Zophobas atratus* Kraatz, 1880 of this species (Yang et al., 2020). As in the mealworms from *T. molitor*, the PS-degrading capabilities of superworms were inhibited by antibiotic suppression of gut microbiota, emphasizing the importance of the gut microbiome in PS degradation (Yang et al., 2020). However, a metagenomic analysis of the superworm gut microbiome, identifying key community members and their inferred functions, is currently missing.

In this study, we aimed to confirm the survival of superworms reared solely on PS, and to investigate changes in the superworm’s gut microbiome in response to this diet. In particular, we compared differences in community compositions and inferred metabolic functions of the gut microbiome from superworms on a standard bran diet, a sole PS diet, and under starvation conditions. We found >95% survival rates among superworms and a marginal weight gain in the PS group compared to starvation conditions. We identified the main microbial players in the gut microbiome during each diet regime, successfully reconstructed microbial genomes and inferred metabolic functions, including enzymes linked to polystyrene and styrene degradation.

## METHODS

### Superworms and polystyrene

Superworms, the larvae of *Zophobas morio*, ranging in size from 25 mm to 40 mm, and superworm pollard (bran) used in this study were purchased from Livefoods Unlimited, Queensland, Australia. Styrofoam was originally a trademarked brand of a light-blue extruded polystyrene (XPS) foam used in building insulation and has since become a generic name for white expanded polystyrene foam (EPA) frequently used in food containers, coffee cups, and packaging material. The styrofoam used in this study was EPA purchased from Polystyrene Products Pty Ltd, QLD, Australia (https://www.polystyreneproducts.com.au/). According to the manufacturer, the styrofoam contained at least 92%-95% polystyrene (PS; CAS No 9003◻53◻6), 4-7% pentane (CAS No 109◻66◻0), an expanding agent added to polystyrene beads, and 1% of the flame retardant Hexabromocyclododecane (CAS No 25637◻99◻4) at the time of manufacture. The styrofoam was stored in our laboratory over several months prior to the feeding trials and was therefore considered containing only trace amounts of pentane, since this agent diffuses to insignificant levels within several weeks after production (Simpson et al., 2020).

### Feeding trials

Initial trials with groups of starving superworms confirmed reports of cannibalism (Maciel-Vergara et al., 2018), which led to our modified experimental design housing the starving control group animals in isolation (**Fig. S1**). While this setup might influence social behavior and could potentially impact the microbiome, we found that isolation is the only effective way to prevent cannibalism under starvation conditions.

A total of 171 superworms were housed in one container for the initial 24 hours following arrival. Subsequently, the 171 superworms were split into three groups with three replicates of 15 superworms each, all of which were fed with wheat bran (certified organic wheat pollard; product code: MWORMPOL; Livefoods Unlimited, QLD, Australia) supplemented with carrots, during an initial one-week acclimatization period (**Fig. S1**). Moisture was controlled with a custom water sprayer to create a fine mist layer on the underside of the superworm container lid. All experiments were carried out at room temperature, which ranged from 20 to 25°C during the experiment. After the acclimatization period, 4 superworms from all three replicates in each group (4*3*3 = 36) were flash-frozen in liquid nitrogen and stored at -80 °C. The remaining 135 superworms underwent the subsequent three-week feeding trial (**Fig. S1**). Thereby each group, consisting of three replicates with 15 superworms each, received a different feed: i) the bran group was fed solely with wheat bran, ii) the polystyrene (PS) group was supplied solely with untreated PS as food source, and iii) the starvation control group received no food supply (**Fig. S1**). The weight of the superworms and all styrofoam blocks was recorded twice a week. After the three-week feeding trial, up to 15 superworms from each group were separated for the pupation trial, and the remaining superworms were flash-frozen in liquid nitrogen and stored at -80 °C for metagenomics.

### Superworm weight statistical analysis

Differences in superworm weight changes between groups were assessed by one-way ANOVA followed by Tukey’s HSD post-hoc test.

### Dissection of the superworms and feces collection

For dissections, the superworms were removed from the -80C storage, placed on their side under a dissection microscope and, while still frozen, cut along the lateral axis from behind the legs to the last abdominal segment. The larvae’s digestive tract was pulled out and midgut and hindgut were removed for DNA extraction. Superworm feces were harvested daily over a one-week time period, in order to accumulate enough material for sequencing analysis, and were stored at RT during that time. After the last collection on day 7 the feces was stored at -80C until DNA extraction.

### DNA extraction and sequencing

The Power Biofilm DNA Isolation Kit (QIAGEN, USA) was used for all microbial DNA extractions. In total, 15 samples were processed, which included 12 superworm gut tissues (three from the acclimatization experiment, and three from each of the three feeding trial groups), a feces sample from the bran group, a feces sample from the PS group, as well as one negative control (200uL water). Genome sequencing was carried out on the Illumina NextSeq 500 platform using Nextera XT libraries, with a targeted sequence allocation of 3Gb per sample.

### Read QC

Quality control of raw Illumina reads (2x 150bp) from all 15 samples was performed with FastQC Version 0.11.9 (https://www.bioinformatics.babraham.ac.uk/projects/fastqc/).

### 16S rRNA / SSU rRNA gene analysis

We characterized the microbial community composition based on small subunit (SSU) rRNA gene abundances with GraftM, a tool that uses gene-specific packages to rapidly identify gene sequences followed by a taxonomic classification derived from placements into a pre-constructed gene tree (Boyd et al., 2018). In brief, SSU rRNA reads were detected and classified using the GraftM package “7.71.silva_v132_alpha1.gpkg” which uses SSU rRNA sequences and the taxonomy-decorated phylogenetic tree based on the 99% nucleotide identity representative OTU set from the SILVA SSU database release 132, Ref NR 99 (https://www.arb-silva.de/). Subsequently, GraftM results were manually curated by removing lineages identified as contamination. In brief, sample contamination is a combination of crosstalk from nearby wells and background contaminants (Minich et al., 2019), and we therefore followed an approach outlined previously (Epstein et al., 2019; Lee et al., 2015), which is based on the presumption that contaminant taxa are expected to have high relative abundances in negative controls but low relative abundances in samples. Therefore, lineages that exhibited a relative abundance of one or more orders of magnitude higher in the negative control compared to any sample were removed. This resulted in the removal of 11 lineages including Cutibacterium, Comamonas and Marinococcus, as well as reads assigned to mitochondria and chloroplasts (**Table S2**). We further evaluated the proportion of eukaryotic host sequences in our samples with graftM (-euk_check) using the package “4.40.2013_08_greengenes_97_OTUs_with_euks.gpkg”.

### Microbial diversity estimates

The Shannon diversity index and the Simpson diversity index were calculated. To negate potential rarefaction biases, we additionally randomly subsampled 500 times and then calculated the average Shannon index.

### Assemblies

We performed genome assembly individually for each sample, as well as combined genome assembly for the three replicates from the same experimental condition using MegaHit v1.1.2 (Li et al., 2015). The individual assemblies were used for gene calling and annotations, whereas the combined assemblies were used for binning.

### Bacterial gene annotation of assemblies - gene-centric analysis

Gene calling of metagenomic assemblies was performed by Prodigal (Hyatt et al., 2010). Genes of metagenome assemblies were then blasted against the Uniprot TrEMBL database (**Fig. S2**) using Diamond’s blastp function (Buchfink et al., 2015). Genes with a bacterial blast top hit were selected and annotated by KOfamScan (Aramaki et al., 2020) for the gene profiling analysis, while eukaryotic genes and genes without blast hits were excluded from the downstream analysis (**Fig. S2**). Gene calling and annotation was performed with enrichM (https://github.com/geronimp/enrichM), which uses HMMS to assign EC, KO, PFAM & TIGRFAM identifiers, and dbCAN to annotate CAZY. To cover assignments not included in the enrichM database, KO IDs were annotated by blasting against Uniprot using Diamond, whereas PFAM and TIGRFAM were searched against their respective databases using hmmsearch. For the functional comparisons, replicates BR3, PS1, and CO1 were excluded from the analysis due to the low gene counts of bacterial genes.

To screen for differentially abundant genes, counts of genes, with a bacterial blast top hit and a KOfamScan annotation (see above) were normalized and transformed with variance stabilized transformation using DESeq2 in R (https://www.rdocumentation.org/packages/DESeq2/versions/1.12.3) from which a Principal component plot (PCA) was created to determine the variation between the communities as well as differential abundances for the top 50 genes. We compared this approach to an alternative method to normalize our data, based on the trimmed mean of M-values (TMM) (Robinson and Oshlack, 2010), using the edgeR package in Bioconductor (Robinson et al., 2010).

To detect enriched KEGG modules, genes with assigned KEGG Orthologs (KO) IDs were rarefied to 8,255 genes, based on the lowest number of genes present in one of our samples, and analyzed with EnrichM classify, which assigns modules based on a matrix of KO annotations (https://github.com/geronimp/enrichM).

### *In silico* PCR amplification

The primers (FW 5’-CGCCAGTTGCTCTGCCATCG-3’, RW 5’-TGCCATGTGG GCGACGCGGC-3’) designed to amplify serine hydrolase (SH) from *Pseudomonas* sp. DSM 50071 (Kim et al., 2020) were used for an *in silico* PCR amplification against *Pseudomonas aeruginosa* on the website http://insilico.ehu.es/PCR/ (Bikandi et al., 2004). The 300bp hit sequence was then blasted (blastX) to obtain SH protein sequences.

### Recovery of metagenome-assembled genomes (MAGs)

Read mapping was performed with BamM v1.7.3 (http://ecogenomics.github.io/BamM/). UniteM (https://github.com/dparks1134/UniteM) and CheckM (Parks et al., 2015) were used for binning and bin quality control, respectively, followed by phylogenetic tree construction and taxonomy assignment performed by gtdb-tk (Chaumeil et al., 2019). Functional analysis was performed with EnrichM (https://github.com/geronimp/enrichM) and KofamScan (Aramaki et al., 2020) as described for the “Bacterial gene annotation of assemblies” above, however without excluding any genes from the analysis.

### Phylogenetic inferences

Phylogenetic trees for the genes of serine hydrolase (SH), StyA, and StyE were inferred from MAFFT (Katoh and Standley, 2013) aligned protein sequences using IQ-TREE 2 (Minh et al., 2020) with the settings “LG+C10+F+G+PMSF” and a starting tree inferred with FastTree 2 (Price et al., 2010).

Phylogenetic trees for metagenome-assembled genomes (MAGs) recovered in this study were inferred based on 122 protein markers aligned with gtdb-tk (Chaumeil et al., 2019) using IQ-TREE 2 (Minh et al., 2020) with the settings “LG+C10+F+G+PMSF” and a starting tree inferred with FastTree 2 (Price et al., 2010).

## RESULTS AND DISCUSSION

### Host survival, behavior and life cycle

Following an initial one-week acclimatization period, we conducted a three-week feeding trial consisting of superworms from three groups; bran reared, polystyrene (PS) reared, and a starvation group (**Fig. S1**). Superworm survival rates in all three groups were above 95% over the entire duration of the feeding trial (**Fig. 1a**). This is in contrast to a previously reported survival rate of ∼70% after three weeks, conducted under similar experimental settings but with over 10 times higher superworm densities (Yang et al., 2020). It is likely that our experimental design prolonged survival by avoiding overcrowding which can lead to aggressive behavior and fatalities among worms, as reported for the common mealworm (Weaver and McFarlane, 1990). Superworms in the PS group readily approached the polystyrene blocks and created narrow burrows by chewing their way into the blocks within the first 24 hours of the experiment (**Fig. 1b**). Worms subjected to the PS diet remained active during the duration of the experiment, although they moved about with a slower speed than worms in the bran group. The starvation group without feed showed the least movement, with prolonged resting periods interspersed by short explorations. The feces changed color from light brown to white pellets in the first 24-48 hours in the PS group, suggesting that the worms had started to consume and egest PS. Dissections of the worm’s digestive tracts revealed that the midgut and hindgut of individuals in the PS group were tightly packed with PS particles (**Fig. S3**). Electron microscopy confirmed that, compared to virgin extruded PS, the PS particles in the superworm guts appeared partially degraded, and showed attached microbial cells (**Fig. S4**). We therefore conclude that superworms were able to ingest PS and pass it through their entire gastrointestinal system, where it came into contact with the gut microbiome.

**Figure 1.**
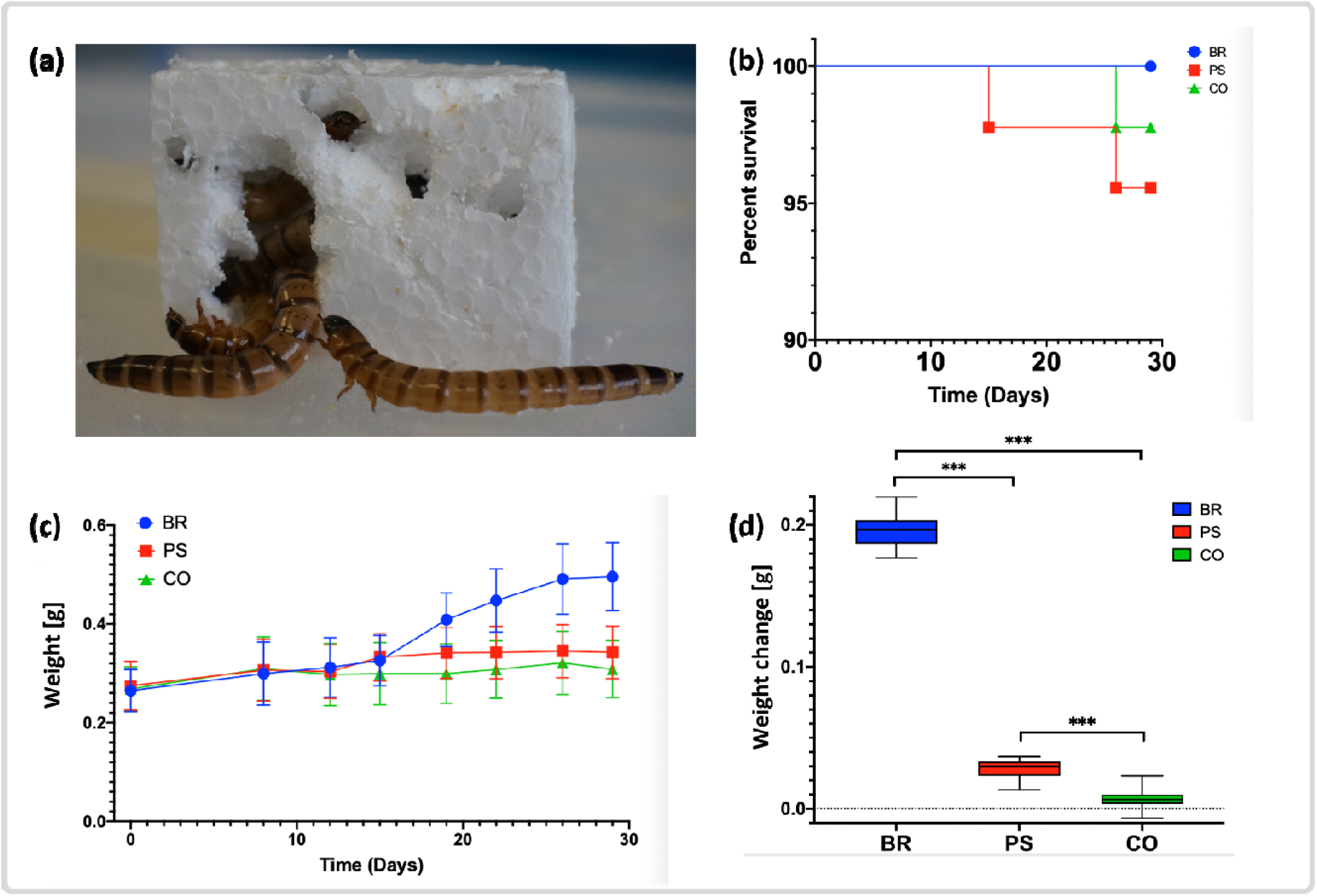
Superworm polystyrene utilization, survival rates, and weight changes. **(a)** Superworm survival rates for the three groups: bran (blue), polystyrene (red), and the control group (green) during the duration of the three-week feeding trial. **(b)** A group of superworms eating their way into a polystyrene block. **(c)** Weight change of the superworms in the three groups. Day 1 to day 7 represents the acclimatization period (ACC). **(d)** Overall weight change of the superworms in each group at the end of the feeding trial. Asterisks indicate that the weight change between BR and PS, BR and CO, and PS and CO was significantly different (**Table S3**). Box plots show the median (bold line), first and third quartile (box outline), 10th and 90th percentile (lower and upper whisker boundaries) and outliers (circles).

The average weight of superworms in the bran group more than doubled during the experiment, leading to significantly heavier worms compared to the PS and the starvation group (**Fig. 1c, d)**. The average superworm weight in the PS group increased slightly during the feeding trial (**Fig. 1c)**, resulting in a marginal but significant weight gain at the end of the feeding trial **(Fig. 1d; Table S3)**. This weight gain and the higher activity compared to the starvation group suggests that worms in the PS group were able to obtain energy from the PS feed, likely in cooperation with their microbial gut communities. This hypothesis is supported by previously reported carbon recovery efficiencies, which indicated that mineralization of ingested PS occurs in the superworm gut (Yang et al., 2020).

To assess the impact of the PS diet on the lifecycle of *Zophobas morio*, we conducted a pupation trial following the three-week feeding experiment (**Fig. S1**). We limited the trial to 10 - 15 worms from each group and observed a pupation rate of 92.9%, 66.7%, 10.0% for the three groups bran, PS, and starvation, respectively (**Fig. S5**). All formed pupae completed the entire pupation phase and emerged as adult beetles. We hypothesize that the marginal weight advantage of worms in the PS group increased the likelihood of a successful pupation nearly 7-fold compared to the starvation group. Next, we analyzed the microbial gut community of the superworms, to identify microbes, encoded pathways, and enzymes potentially involved in PS degradation.

### Microbial community composition

Mid- and hindguts of superworms (**Fig. S3**) were dissected and, together with two fecal samples, used for DNA extraction and metagenomic sequencing (*see Methods*). The obtained shotgun reads were mined for small subunit (SSU) rRNA gene sequences to omit insect host genes (**Fig. S6**) and to characterize prokaryotic microbial gut communities. On average, the microbial diversity decreased from bran to PS and further from PS to starvation conditions (**Table S4**), suggesting that PS is a poorer diet compared to bran, but still supports a more diverse community compared to starvation conditions. This conclusion aligns with previous reports demonstrating that microbiome diversity is positively correlated with diet diversity (Heiman and Greenway, 2016), a trend that has also been confirmed for insect larvae (Krams et al., 2017; Priya et al., 2012). The lowest microbial diversity, observed in the starvation group, indicates a lack of nutrients or even the onset of a disease. The latter assumption is supported by the detection of pathogenic bacteria (see below) and reports that a loss of gastrointestinal species diversity is a common finding in several vertebrate disease states (Heiman and Greenway, 2016). This correlation might translate to insect gut microbiomes, since greater microbial diversity has been associated with health benefits for insect hosts (Mattila et al., 2012).

Overall, the gut microbiome of the baseline acclimation, bran, PS, and starvation group was dominated by the two bacterial phyla Firmicutes and Proteobacteria with maximum relative abundances of 80.6% and 67.8%, respectively (**Fig. 2; Table S5, S6**). Other phyla, including Tenericutes, Bacteroidetes, and Actinobacteria were also present, although at lower relative abundances with a maximum of 15.9%, 15.1%, and 11.5%, respectively (**Table S6, Fig. 2**). This observed superworm microbial gut community is similar to microbiomes from common mealworms (*T. molitor*), with studies reporting Firmicutes, Proteobacteria but also Tenericutes and Actinobacteria as dominant phyla (Jung et al., 2014; Wang and Zhang, 2015). Interestingly, no archaeal SSU rRNA sequences were detected in our superworm guts, similar to findings from a previous analysis of mealworms (Urbanek et al., 2020). This likely indicates that Archaea are only present in very low abundances below detection limits or are not part of mealworm and superworm gut microbiomes.

**Figure 2.**
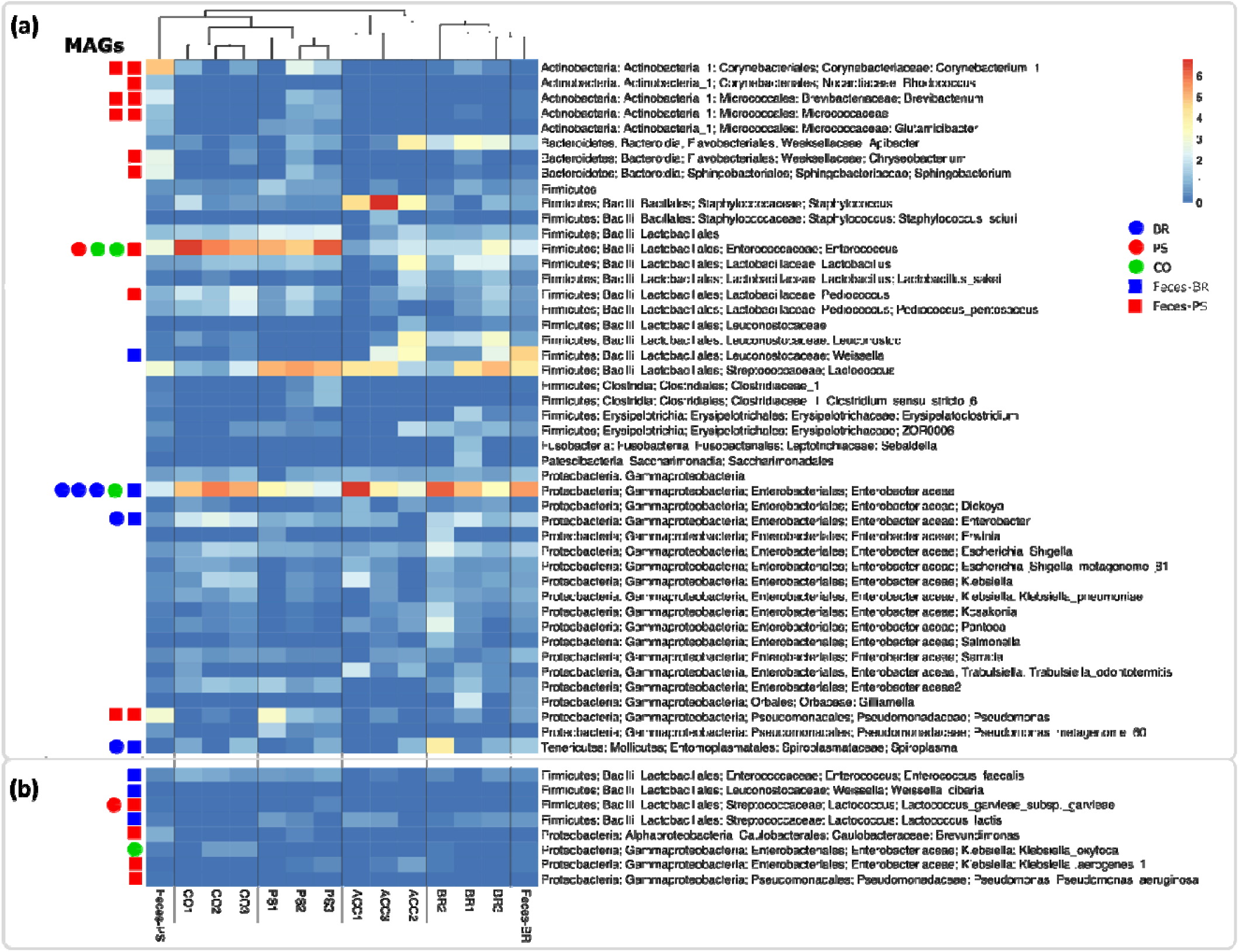
Abundant bacterial taxa in the superworm gut microbiome. The profile is based on SSU rRNA short read alignments against the Silva r132_99 database). (**a**) Taxonomic lineages with a relative abundance >1% were included in the heatmap. Abbreviations: CO1-3 (replicates of the starvation control group without feed), PS1-3 (replicates of the polystyrene group), ACC1-3 (replicates of the acclimation baseline group sampled before the feeding trial), BR1-3 (replicates of the wheat bran group), Feces-PS (polystyrene feces), Feces-BR (bran feces). Metagenome-assembled genomes (MAGs) recovered from our samples are indicated with a circle (feeding groups) or a square (polystyrene and bran feces samples) next to the assigned lineages. (**b**) Eight lineages with a relative abundance <1% from which we recovered metagenome-assembled genomes (MAGs).

Among the most abundant taxa in the bran group, were facultative anaerobes from the Gammaproteobacteria family Enterobacteriaceae, including the genera *Enterobacter, Escherichia/ Shigella, Erwinia, Klebsiella* and *Dickeya* (**Fig. 2**). These taxa have been reported from insect guts previously (Jurkevitch, 2011) and some lineages including *Enterobacter, Klebsiella* and *Erwinia* are known for mutualistic relationships with their insect hosts (Dillon and Dillon, 2004; Gupta and Nair, 2020; Gurung et al., 2019; Jurkevitch, 2011). Other genera such as *Dickeya* contain mainly plant pathogens (Samson et al., 2005) with a potential to kill insect hosts (Costechareyre et al., 2012). Overall, the bran microbiome did not change significantly compared to the baseline group (**Fig. 2**), suggesting that the worms became well acclimatized before the feeding trials commenced. The only exception was an uncultured species in the genus *Staphylococcus* which showed significantly higher relative abundance in the acclimation group compared to all other groups (**Fig. 2; Fig. 3a, b, c**). Further abundant taxa in the bran group were lactic acid bacteria from the class Bacilli including the genera *Lactoccocus, Weisella*, and *Leuconostoc*, which contain species with reported probiotic effects (Fusco et al., 2015; Shuttleworth et al., 2019). In particular, *Weisella, Leuconostoc*, and *Lactoccocus* were significantly enriched in the bran group compared to the PS and starvation group, respectively (**Fig. 3d,e**), indicating that these taxa might benefit from a fiber-rich bran diet.

**Figure 3.**
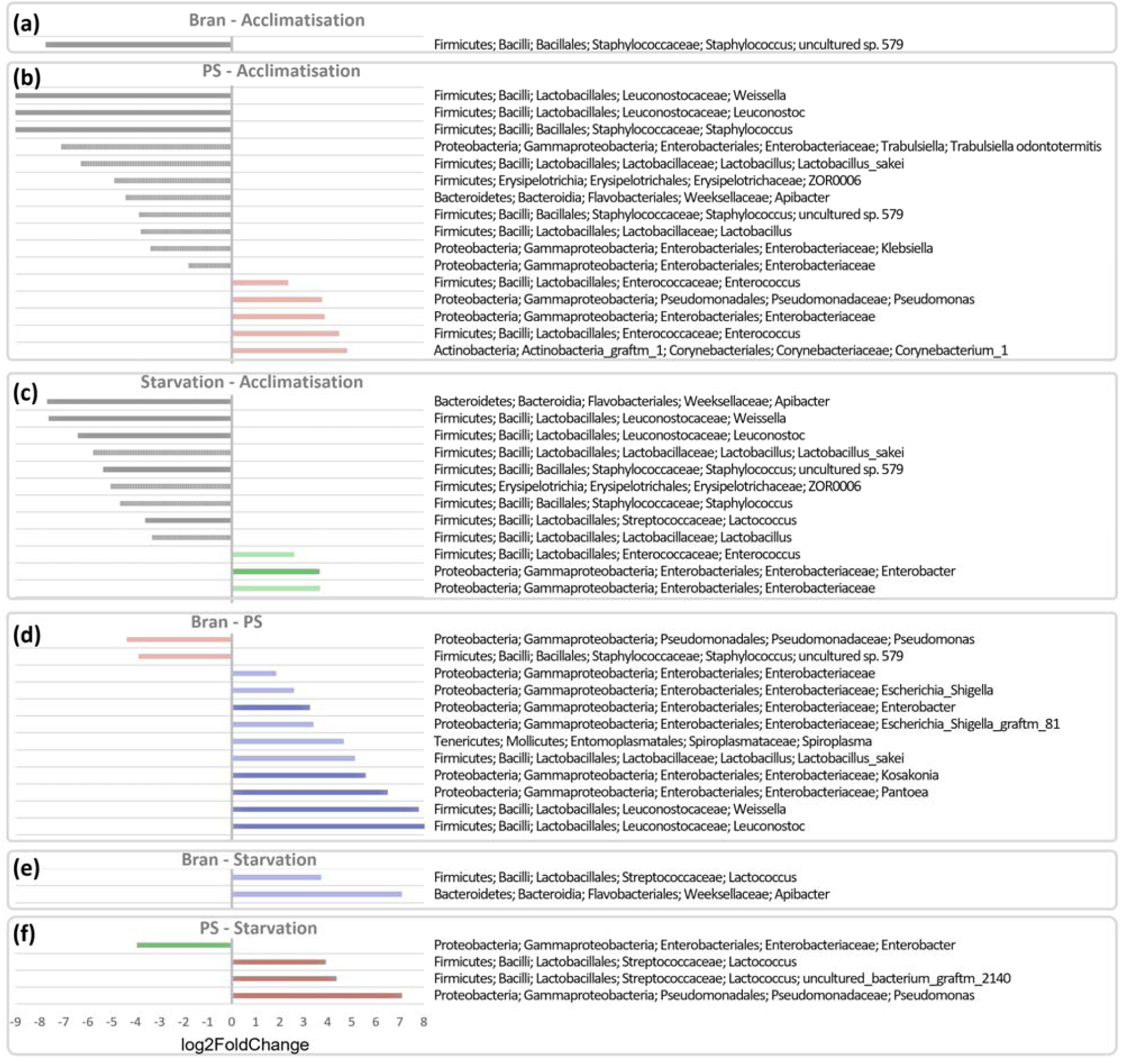
Gut microbiome lineages with significant differences in abundances between feeding trial groups. Estimated log2fold changes with an adjusted p-value of < 0.1 (light bars) and < 0.05 (dark bars) comparing the feeding trials: **a)** bran vs acclimatization, **b)** polystyrene vs acclimatization, **c)** starvation vs acclimatization, **d)** bran vs polystyrene, **e)** bran vs starvation, and **f)** polystyrene vs starvation. Abbreviations: polystyrene (PS). Color code: bran (blue), polystyrene (red), and the starvation control group (green).

The microbiome of the PS group superworms differed from the bran group by a lower relative abundance of the family Enterobacteriaceae (**Fig. 2**). In particular, the Enterobacteriaceae genera *Kosakonia, Pantoea, Weisella, Leuconostoc*, and *Enterobacter* had significantly lower abundances compared to the bran group (**Fig. 3d**). The PS group was further characterized by higher relative abundances of the Bacilli genera *Lactococcus* and *Enterococcus*, whereby the latter genus was also elevated in the starvation group (**Fig. 2**). The presence of *Enterococcus*, a genus of facultative anaerobes (Dubin and Pamer, 2014), in the superworm gut is not surprising since several *Enterococcus* species were isolated from gastrointestinal tracts of animals, including a variety of insects (Lebreton et al., 2014). Members of this genus have also been associated with plastic substrates. Multiple *Enterococcus* species were found to survive over 90 days on polyester and polyethylene fabrics in hospitals (Neely and Maley, 2000), and elevated levels of *E. avium* were reported from mealworms reared on a diet of extruded PS (Urbanek et al., 2020). Our data confirms the association of *E. avium* with a PS diet, since we detected this species in the PS but not in the bran and starvation groups (**Table S7**), although with low relative abundances (<0.2%). However, given the large phylogenetic and metabolic diversity of the genus *Enterococcus* (Dubin and Pamer, 2014), it is possible that some taxa act as pathobionts in the insect gut, similar to reports from mammals (Balish and Warner, 2002).

In addition to superworm gut samples, we analyzed a fecal sample from the PS and the bran group, respectively. While the bran fecal sample clustered with the bran group, the PS fecal sample was distinct from all other samples. The dominating lineage was *Corynebacterium*, a genus that was also moderately abundant in the PS group (**Fig. 2**). The genus *Corynebacterium* comprises mostly aerobic bacteria associated with a range of host organisms, including humans (Wallen et al., 2020). *Corynebacterium*, as defined in NCBI, is divided into two genera in the SILVA database, named *Corynebacterium* and *Corynebacterium_1*. The latter genus is present in our samples **(Fig. 2)** and has been reported as an opportunistic pathogen linked to dysbiosis in the gut microbiome of patients with Parkinson’s disease (Wallen et al., 2020).

The starvation group gut microbiome shared the high relative abundance of *Enterococcus* with the PS group but had a significantly lower level of *Lactococcus* and a significantly higher level of *Enterobacter* compared to the PS group (**Fig. 2; Fig. 3**). The elevated presence of the pathogen *Klebsiella oxytoca* **(Fig. 2**), an emerging human pathogen causing hospital-acquired infection (Singh et al., 2016), supports previous findings of an elevated *K. oxytoca* presence in starved insect larvae, based on experiments with the common mealworm *Tenebrio molitor* (Urbanek et al., 2020).

In conclusion, we hypothesize that higher relative abundances of *Enterococcus* in the PS and starvation groups, of *K. oxytoca* in the starvation group, and of *Corynebacterium_1* in the PS feces, indicate that a PS diet and starvation conditions result in poor gut health and potentially induce dysbiosis in the superworm gut. Future studies combining microbial community profiling and gut histomorphology are necessary to test this hypothesis.

### Microbial taxa associated with polystyrene degradation

Over the last two decades, a range of bacterial strains have been enriched or isolated with PS as the sole carbon source, and most were identified at the genus or species level (**Table S8**). Based on our SSU assignments, we detected six of these genera (*Bacillus, Brevundimonas, Microbacterium, Pseudomonas, Sphingobacterium*, and *Streptococcus*) with higher relative abundances in the PS group compared to all other groups (**Table S8**). In particular, *Pseudomonas* had significantly higher relative abundances in the PS group compared to the bran group (**Fig. 3d**). Various species from this genus have been previously isolated with PS as the sole carbon source (**Table S1**), and we detected two, *Pseudomonas aeruginosa* and *P. putida*, solely in the PS feces, albeit with low relative abundances below 0.02% (**Table S8**). Further PS degrading taxa, classified at the species level (**Table S1**), detected in our samples include *Brevundimonas diminuta* uniquely detected in the PS group, and *Rhodococcus ruber exclusively* detected in the PS feces (**Table S8**). Interestingly, the genus *Kosakonia*, reported to have a strong association with PS and polyethylene diets in the common mealworm (Brandon et al., 2018), had a significantly lower abundance in the PS group compared to the bran group (**Fig. 3d**). This suggests that different *Kosakonia* species or strains could be present in the gut of superworms and mealworms, respectively. Future deep metagenomic sequencing might be able to provide the necessary taxonomic resolution to resolve this question.

### Functional community profiles

To assess changes in the functional potential of gut microbiomes across samples, we assigned genes to bacterial KEGG Ortholog (KO) identifiers (**Table S9**). Overall, the encoded microbiome functions differed considerably among bacterial genes from the bran, PS and the starvation groups, while clustering according to diet (**Fig. 4a**). The PS group was characterized by the absence or incompleteness of several biosynthesis modules, including amino acid, lipopolysaccharide and GABA biosynthesis, as well as a lack of antimicrobial peptides, drug resistance, transport and two-component regulatory systems (**Fig. 4b**). Modules enriched in the PS group include transporters such as the putative aldouronate transport system (**Fig. 4b**), which has been reported to facilitate the intracellular conversion of aldouronates following extracellular depolymerisation of hemicellulose in bacteria (Chow et al., 2007). However, whether these transporters aid in the import of depolymerized PS remains to be determined.

**Figure 4.**
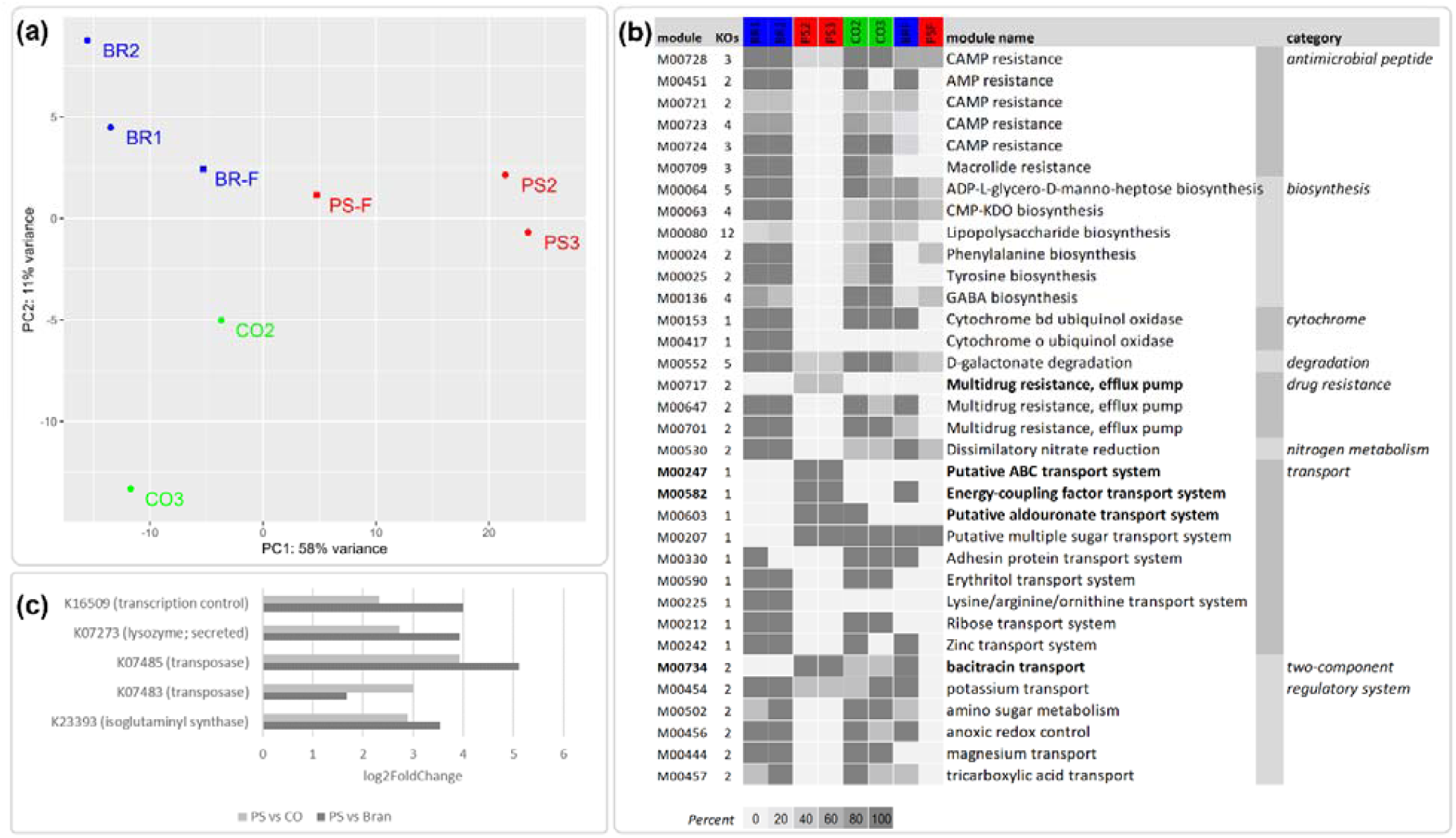
Functional profiles of superworm microbiomes. **(a)** Principal component analysis (PCA) of genes assigned to bacterial KEGG Orthologs (KO identifiers), showing a clear separation of all three feeding trials, including bran (BR; blue dots), polystyrene (PS; red dots); starvation control group (CO; green dots), as well as polystyrene feces (PSF; red squares) and bran feces (BRF; blue squares). **(b)** KEGG modules enriched and depleted in the polystyrene (PS) group based on rarefied gene counts. Shown are the KEGG modules, the number of KOs per modules, and the module completeness (0-100%) reported for the bran samples (BR1, BR2), polystyrene samples (PS2, PS3), the starvation control samples (CO2, CO3). **(c)** Differential abundant genes, with KO identifiers, enriched in the PS group compared to the starvation control (CO) and bran group, respectively. Note, that gene counts were rarefied to 8,255 for the module comparison in (b).

A small subset of genes assigned to functional homologs were differentially more abundant in the PS group compared to the bran and the starvation group (**Fig. 4c**), indicating a possible association with a PS diet. Two genes were identified as bacterial transposases (**Fig. 4c; Table S10**). Transposases are known to facilitate gene duplications and genomic rearrangements in bacteria, and are considered to be means of genome adaptations to survive stressful situations following environmental changes (Casacuberta and González, 2013). The unbalanced diet in the PS group could represent such challenging conditions for the bacterial gut community, resulting in higher abundances of transposases as a strategy to facilitate rapid adaptation to cope with the stressful environment (Li et al., 2014). Genes enriched in the PS group also suggest that the PS microbiome has a greater potential for membrane restructuring. This conclusion is supported by enriched genes encoding the lysosome M1 (GH25 family, K07273), an enzyme involved in remodeling peptidoglycan and in cell lysis for the dissemination of phage progeny (Vollmer et al., 2008), and isoglutaminyl synthase (K23393, murT) involved in peptidoglycan biosynthesis (Figueiredo et al., 2012). Whether this increased membrane plasticity is another stress response needs to be determined. However, the transcriptional regulator protein spxA (K16509), which is responsible for transcriptional control during oxidative stress and may cause genome-wide changes in expression patterns (Zuber, 2004) was also enriched in the PS group, indicating that increased oxygen exposure might be a stress contributing factor. We screened for aerobic and anaerobic modules across all samples to investigate if the nearly anoxic conditions, which exist in insect guts (Johnson and V. Barbehenn, 2000), might have been influenced by the PS diet. Our results showed an increased presence of encoded aerobic functions in the PS group and the PS feces, compared to all other samples (**Table S11**). This supports our conclusion of deteriorating gut conditions in the PS group (see “*Microbial community composition*”), since a healthy gut epithelial is expected to consume oxygen, generating a state of hypoxia in humans (Valdes et al., 2018) and likely also in insects (Tegtmeier et al., 2016).

Furthermore, we hypothesized that a shift in diet from bran, i.e. wheat pollard that is a by-product of the flour milling of grain and has a high fiber content (Stevenson et al., 2012), to PS will lead to a fiber deprived gut microbiome. Such a microbiome can degrade the colonic mucus barrier and enhance pathogen susceptibility, as shown in mouse models (Desai et al., 2016). Indeed, a functional characterization of encoded carbohydrate active enzymes (CAZy) revealed differences between samples (**Fig. S7**) but did not detect enriched or depleted fiber or mucin targeting CAZy genes, likely due to the fact that such changes commonly occur on a transcriptional level (Desai et al., 2016).

### Detection of potential polystyrene and styrene degrading enzymes

The lack of characterized enzymes for the initial step in microbial PS degradation, the breakdown of the insoluble PS polymer into styrene monomers, has led to the conclusion that the environmental degradation of this synthetic polymer depends initially on abiotic factors. UV irradiation and mechanical forces, such as wave action, wind, climate, or presumably the shredding and ingesting of PS by superworms, attack the PS backbone that contains only carbon-carbon bonds and has no “hydrolyzable” groups. This abiotic stress causes the polymer to age, that is, to react with small environmental molecules, typically O_2_ or H_2_O, resulting in the formation of carbonyl groups (Audouin et al., 1994) which are considered more accessible to enzymatic degradation (Albertsson and Karlsson, 1990; Fontanella et al., 2010; Koutny et al., 2006). However, microbial pathways to age PS polymers seem plausible and could utilize hydroxylases for oxygen insertion (Lewis et al., 2011) and subsequently dehydrogenases to form carbonyl groups (**Fig. 5**). This process could be catalyzed by extracellular enzymes, similar to fungal extracellular cellobiose dehydrogenases (Ma et al., 2017). Subsequently, enzymes with catalytic activities similar to phenolic acid decarboxylases (Frank et al., 2012), lipases (Mohan et al., 2016), serine hydrolase (Kim et al., 2020), or the recently characterized PET degrading enzyme PETase (Chen et al., 2018) could facilitate a further PS breakdown by targeting carbonyl groups. Currently, the best evidence for an enzymatic carbonyl group-based PS degradation comes from an analysis of the aforementioned *serine hydrolase* (SH) gene, reported from a *Pseudomonas aeruginosa* strain isolated from a superworm gut under oxygen free conditions (Kim et al., 2020). The authors report that qPCR-based SH expression levels were elevated when growing the *Pseudomonas* strain on a nutrient-limited medium with added PS. Furthermore, PS degradation did not occur when the SH was blocked with an inhibitor, leading the authors to conclude that SH is involved in the breakdown of PS by targeting carbonyl groups, likely resulting in styrene dimers or monomers (Kim et al., 2020). Since the authors did not sequence the reported SH gene (Kim et al., 2020), we performed an *in silico* PCR, followed by a blastp detection of this gene in our dataset. Among the feeding trials, we recovered SH homologues solely from the PS group, however both PS and bran feces assemblies contained copies of this gene as well (**Table S12**). Phylogenetic inferences revealed that the SH genes recovered in our study clustered with the homologue from *P. aeruginosa*, indicating that the encoded superworm microbiome SH proteins can facilitate PS breakdown (**Fig. S8**). However, further culture-based experiments, including transcriptomics and gene knockout trials, are necessary to clarify the function of SH and other genes in the PS degradation pathway of this and related *P. aeruginosa* strains present in the superworm gut microbiome.

**Figure 5.**
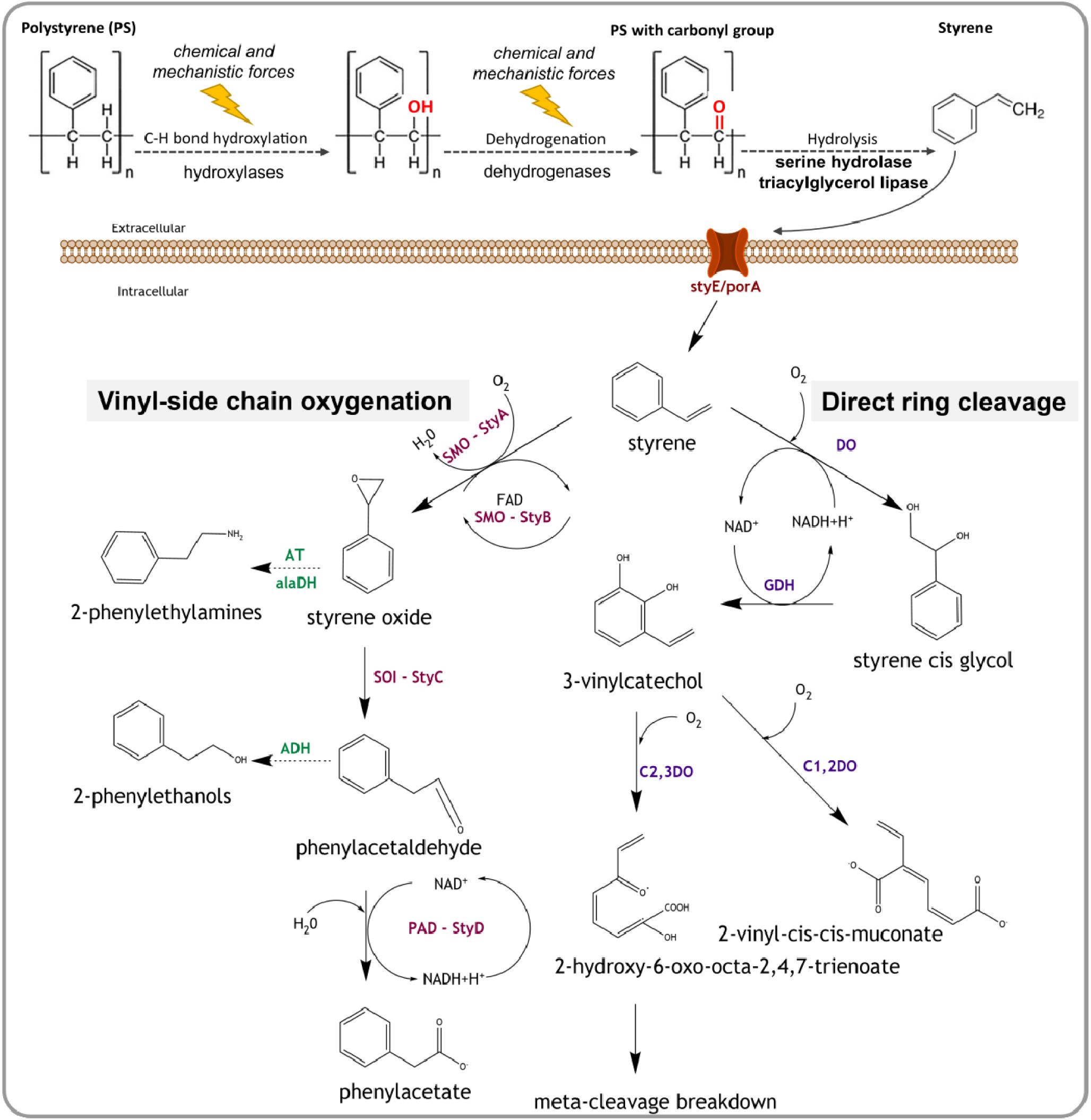
Potential pathways and enzymes of bacterial polystyrene and styrene degradation. Initially, abiotic chemical and physical factors, and potentially also extracellular enzymes such as hydroxylases and dehydrogenases, attack the polystyrene polymer creating carbonyl groups. Next, extracellular enzymes such as serine hydrolase and triacylglycerol lipase degrade the polymer to styrene monomers. A putative transmembrane protein StyE/PorA can import styrene monomers into the cell, where it is degraded via the vinyl side chain oxygenation or the direct ring cleavage pathway. In the **vinyl side chain oxygenation pathway**, styrene is broken down to phenylacetate using enzymes of the styABCD operon: styrene monooxygenase (StyA/SMO), styrene monooxygenase reductase component (StyB), styrene oxide isomerase (StyC/SOI), and phenylacetadehyde dehydrogenase (StyD/PAD/FeaB/ TynC). Styrene can also form 2-phenylethylamines and 2-phenylethanols as intermediates, through modifications of vinyl side chain pathway by the enzymes aminotransferase (AT), alanine dehydrogenase (alaDH), and alcohol dehydrogenase (ADH). **The direct ring cleavage pathway** converts styrene into krebs cycle intermediates via meta cleavage breakdown or to muconic acid. Involved key enzymes are styrene dioxygenase (DO), 2,3-dihydrodiol dehydrogenase/cis glycol dehydrogenase (GDH), catechol-1,2-dioxygenase (C1,2DO), and catechol-2,3-dioxygenase (C2,3DO). Dashed arrows indicate hypothetical reactions.

Another group of hydrolases with a potential function in polymer degradation are esterases. In particular, lipases, a subclass of esterases, have been experimentally verified to perform surface hydrolysis of synthetic polymers, including polyethylene terephthalate (PET) fabrics and films (Eberl et al., 2009). Furthermore, triacylglycerol lipase (EC 3.1.1.3), as well two esterase carboxylesterases (EC 3.1.1.1) and cutinases (EC 3.1.1.74), have been identified as probable enzymes for the enzymatic hydrolysis of difficult to degrade aliphatic-co-aromatic polyesters (Kawai et al., 2019). An unspecified lipase has also been reported from a high impact polystyrene (HIPS) degrading *Bacillus* spp. (Mohan et al., 2016). Based on this data, we screened the genes recovered from all samples for lipase, carboxylesterase, and cutinase homologues. The only esterases’ genes detected were homologues encoding ***triacylglycerol lipase***, which we found exclusively in the PS group and in the PS feces (**Table S13**). We therefore hypothesize that this lipase homologue could be involved in polystyrene degradation by acting on the carbonyl group of the partially degraded PS, next to its traditional function of attacking the ester carbonyl of fatty acids. Furthermore, lipases are known to act on medium to long-chain polymers with more than ten carbon atoms (W. Yang et al., 2015), which makes these enzymes potentially more suitable for the degradation of partially degraded synthetic polymers; however, experimental verification is needed to test this hypothesis.

Once PS is degraded to styrene, these monomers can be further broken down by a variety of microbial pathways, owing to the fact that styrene occurs naturally in plants and fungi (Warhurst and Fewson, 1994). Two main pathways have been described to facilitate this breakdown under aerobic conditions, the direct ring cleavage and the vinyl side chain oxygenation (Bestetti et al., 2004; Tischler, 2015; Warhurst et al., 1994). Initially, the side chain oxygenation was considered to be more common, but a recent study reported that just 14 out of 87 styrene utilizing strains encode this pathway, indicating that the direct ring cleavage, or a modified version of this pathway, is more widespread than initially thought (Oelschlägel et al., 2014). Direct ring cleavage includes an initial dihydroxylation of the aromatic ring to form styrene cis-glycol, followed by the actual ring cleavage (**Fig. 5**) and further degradation via the ortho or meta cleavage pathway of aromatic compounds, to derive central intermediates such as pyruvate, acrylic acid and acetaldehyde (Patrauchan et al., 2008; Tischler, 2015; Tischler and Kaschabek, 2012; Warhurst et al., 1994). Key enzymes in the direct ring cleavage include styrene dioxygenase, which catalyses the first step in this pathway but currently lacks gene sequences, and glycol dehydrogenase (GDH) mediating the subsequent conversion to vinylcatechol (**Fig. 5**). We detected GDH homologues in nearly all of our samples (**Table S13**), indicating that this dehydrogenase is involved in many aspects of aromatic compound metabolism, and probably not restricted to styrene degradation.

In the second pathway, the vinyl side chain oxygenation, styrene is degraded to styrene oxide via monooxygenases and eventually to phenylacetic acid (**Fig. 5**), leaving the aromatic ring intact (Beltrametti et al., 1997; Bestetti et al., 2004; Itoh et al., 1997; O’Connor et al., 1995). We detected genes for a complete styrene monooxygenase *styA/styB* (**Fig. 5**) solely in the PS feces group (**Table S13**), likely because feces represent an aerobic environment. To assess the evolutionary history of styrene monooxygenase, we inferred the phylogeny of *styA* genes and found that, while the overall gene phylogeny aligns with taxonomic classifications, there is evidence for considerable horizontal gene transfer (**Figure S9**). StyA genes recovered from the superworm feces were grouped with experimentally verified homologues, e.g. from *Pseudomonas fluorescens* (Beltrametti et al., 1997), suggesting that they indeed encode functional styrene degrading enzymes. Furthermore, homologues of *styB*, without the presence of *styA*, were detected in the bran and starvation group, indicating that these gene homologues encode enzymes functioning in pathways unrelated to styrene degradation. Also, *styD* genes, encoding phenylacetaldehyde dehydrogenase (**Fig. 5**), were present in high numbers in all feeding trial groups (**Table S13**), confirming previous findings that *styD* is strongly homologous to a range of prokaryotic aldehyde dehydrogenases (Beltrametti et al., 1997). Distant homologues of *styE* genes, encoding an outer membrane protein reported to be involved in styrene transport in *Pseudomonas putida* (Mooney et al., 2006), were detected in PS feces and bran assemblies (**Table S14**). Phylogenetic inferences clustered the superworm microbiome *styE* homologues with genes for toluene catabolism membrane proteins (Kahng et al., 2000) in a sister clade to a clade containing the originally described *styE* gene (**Fig. S10**). Therefore, *styE* might be involved in styrene transport in multiple genera, however, functional validations are required to determine the role of the *styE* homologues recovered from the superworm gut.

Anaerobic transformation of styrene has been observed in microbial consortia isolated from sewage sludge, and degradation routes via phenylacetaldehyde and phenylacetate, similar to the side chain oxidation, or via ethylphenol have been proposed (Grbić-Galić et al., 1990; Tischler, 2015). However, the involved enzymes remain to be characterized, and hence we could not assess their presence in the superworm gut microbiome.

### Linking Phylogeny and function

To associate the inferred metabolic capabilities of the superworm gut microbiome with the taxa observed in our community profiles, we binned metagenome-assembled genomes (MAGs) from assemblies of all samples. We recovered 34 medium-to-high-quality MAGs with an average estimated completeness of 90.0% (±9.7%) and a mean estimated contamination of 1.3% (±0.9%) (**Table 1**). In addition, we also included one low-quality draft MAG (48.8% estimated completeness) harboring potential styrene degradation genes (**Table 1**). Taxonomic classification assigned the MAGs to 21 genera in four phyla: Proteobacteria, Firmicutes, Actinobacteria, and Bacteroidota (**Table 1, Fig. S11**). MAGs assigned to the phylum Bacteroidota, orders Flavobacteriales and Shingobacteriales, and to the phylum Actinobacteria were solely recovered from PS feces (**Fig. S11**), which was expected given the higher relative abundances of these lineages in the PS feces community profile (**Fig. 2**).

**Table 1.** MAGs | Metagenome-assembled genomes (MAGs) statistics. MAGs recovered from the polystyrene (ps), bran (bran), and starvation (nf) group, as well as from the polystyrene feces (feces_ps) and the bran feces (feces_b). Taxonomic classifications were carried out with GTDB-TK (R202) and estimated completeness, estimated contamination, and strain heterogeneity were assessed with CheckM. GTDB assigned genera and species were screened against the ABSA International’s Risk Group Database (https://my.absa.org/tiki-index.php?page=Riskgroups) to identify potential pathogens (Pathogen). Taxonomic classifications (GTDB-R202) are provided as GTDB tax string. Abbreviations: Estimated completeness (by CheckM) = Comp; Estimated contamination (by CheckM) = Cont; Estimated strain heterogeneity (by CheckM) = Strain. The genome quality (Q) is calculated as Comp -5x Cont. The only low-quality MAG included in our analysis is highlighted in italic.

| MAG ID | gtdbtk-r202 tax string | Comp | Cont | Strain | Q |
| --- | --- | --- | --- | --- | --- |
| ps_bin.7 | d__Bacteria;p__Firmicutes;c__Bacilli;o__Lactobacillales;f__Streptococcaceae;g__Lactococcus;s__Lactococcus garvieae_A | 86.11 | 0 | 0 | 86.11 |
| ps_bin.4 | d__Bacteria;p__Proteobacteria;c__Gammaproteobacteria;o__Pseudomonadales;f__Pseudomonadaceae;g__Pseudomonas;s__Pseudomonas aeruginosa | 48.8 | 0.56 | 0 | 46 |
| ps_bin.3 | d__Bacteria;p__Firmicutes;c__Bacilli;o__Lactobacillales;f__Enterococcaceae;g__Enterococcus_C;s__Enterococcus_Csp009933135 | 85.26 | 1.62 | 28.57 | 77.16 |
| nf_max107.003 | d__Bacteria;p__Proteobacteria;c__Gammaproteobacteria;o__Enterobacterales;f__Enterobacteriaceae;g__LSJC7;s__LSJC7sp005671395 | 99.03 | 0.74 | 0 | 95.33 |
| nf_bin.6 | d__Bacteria;p__Proteobacteria;c__Gammaproteobacteria;o__Enterobacterales;f__Enterobacteriaceae;g__Klebsiella_A;s__Klebsiella_A oxytoca | 98.2 | 0.19 | 50 | 97.25 |
| nf_bin.5 | d__Bacteria;p__Firmicutes;c__Bacilli;o__Lactobacillales;f__Enterococcaceae;g__Enterococcus_C;s__Enterococcus_Csp009933135 | 74.94 | 2.69 | 50 | 61.49 |
| nf_bin.1 | d__Bacteria;p__Firmicutes;c__Bacilli;o__Lactobacillales;f__Enterococcaceae;g__Enterococcus_D;s__ | 96.62 | 2.51 | 7.69 | 84.07 |
| feces_ps_gm2_bin_9 | d__Bacteria;p__Proteobacteria;c__Gammaproteobacteria;o__Enterobacterales;f__Enterobacteriaceae;g__Klebsiella;s__Klebsiella aerogenes | 77.78 | 3.11 | 50 | 62.23 |
| feces_ps_gm2_bin_7 | d__Bacteria;p__Proteobacteria;c__Gammaproteobacteria;o__Pseudomonadales;f__Pseudomonadaceae;g__Pseudomonas_E;s__Pseudomonas_E qingdaonensis | 97.88 | 0.88 | 28.57 | 93.48 |
| feces_ps_gm2_bin_6 | d__Bacteria;p__Proteobacteria;c__Gammaproteobacteria;o__Pseudomonadales;f__Pseudomonadaceae;g__Pseudomonas_E;s__Pseudomonas_E sp001320525 | 93.56 | 1.31 | 25 | 87.01 |
| feces_ps_bin.9 | d__Bacteria;p__Actinobacteriota;c__Actinomycetia;o__Actinomycetales;f__Micrococcaceae;g__Galactobacter;s__ | 90.13 | 1.76 | 0 | 81.33 |
| feces_ps_bin.7 | d__Bacteria;p__Bacteroidota;c__Bacteroidia;o__Flavobacteriales;f__Weeksellaceae;g__Chryseobacterium;s__ | 91.91 | 0 | 0 | 91.91 |
| feces_ps_bin.6 | d__Bacteria;p__Actinobacteriota;c__Actinomycetia;o__Mycobacteriales;f__Mycobacteriaceae;g__Rhodococcus;s__Rhodococcus hoagii | 91.51 | 2.95 | 3.23 | 76.76 |
| feces_ps_bin.5 | d__Bacteria;p__Actinobacteriota;c__Actinomycetia;o__Actinomycetales;f__Brevibacteriaceae;g__Brevibacterium;s__ | 97.88 | 1.35 | 33.33 | 91.13 |
| feces_ps_bin.3 | d__Bacteria;p__Proteobacteria;c__Gammaproteobacteria;o__Pseudomonadales;f__Pseudomonadaceae;g__Pseudomonas;s__Pseudomonas aeruginosa | 85.28 | 1.38 | 11.11 | 78.38 |
| feces_ps_bin.21 | d__Bacteria;p__Firmicutes;c__Bacilli;o__Lactobacillales;f__Streptococcaceae;g__Lactococcus;s__Lactococcus garvieae_A | 98.09 | 0.2 | 50 | 97.09 |
| feces_ps_bin.2 | d__Bacteria;p__Actinobacteriota;c__Actinomycetia;o__Myco<br>bacteriales;f__Mycobacteriaceae;g__Corynebacterium;s__ | 98.12 | 0.83 | 0 | 93.97 |
| feces_ps_bin.19 | d__Bacteria;p__Firmicutes;c__Bacilli;o__Lactobacillales;f__E<br>nterococcaceae;g__Enterococcus_C;s__Enterococcus_C<br>sp009933135 | 90.81 | 1.98 | 15.38 | 80.91 |
| feces_ps_bin.16 | d__Bacteria;p__Proteobacteria;c__Alphaproteobacteria;o__C<br>aulobacteriales;f__Caulobacteraceae;g__Brevundimonas;s__ | 83.17 | 0.35 | 50 | 81.42 |
| feces_ps_bin.15 | d__Bacteria;p__Actinobacteriota;c__Actinomycetia;o__Actin<br>omycetales;f__Brevibacteriaceae;g__Brevibacterium;s__ | 89.53 | 1.94 | 25 | 79.83 |
| feces_ps_bin.14 | d__Bacteria;p__Actinobacteriota;c__Actinomycetia;o__Myco<br>bacteriales;f__Mycobacteriaceae;g__Corynebacterium;s__ | 98.42 | 0.56 | 0 | 95.62 |
| feces_ps_bin.12 | d__Bacteria;p__Actinobacteriota;c__Actinomycetia;o__Actin<br>omycetales;f__Micrococcaceae;g__Galactobacter;s__ | 95.4 | 0.46 | 0 | 93.1 |
| feces_ps_bin.10 | d__Bacteria;p__Bacteroidota;c__Bacteroidia;o__Sphingobact<br>eriales;f__Sphingobacteriaceae;g__Sphingobacterium;s__ | 97.86 | 1.46 | 0 | 90.56 |
| feces_ps_bin.1 | d__Bacteria;p__Firmicutes;c__Bacilli;o__Lactobacillales;f__La<br>ctobacillaceae;g__Pediococcus;s__ | 73.2 | 1.81 | 16.67 | 64.15 |
| feces_b_bin.9 | d__Bacteria;p__Proteobacteria;c__Gammaproteobacteria;o__<br>_Enterobacterales;f__Enterobacteriaceae;g__LSJC7;s__LSJC7<br>sp005671395 | 78.28 | 1.82 | 27.78 | 69.18 |
| feces_b_bin.8 | d__Bacteria;p__Proteobacteria;c__Gammaproteobacteria;o__<br>_Enterobacterales;f__Enterobacteriaceae;g__Enterobacter;s__<br>_Enterobacter oligotrophicus | 62.79 | 0 | 0 | 62.79 |
| feces_b_bin.5 | d__Bacteria;p__Firmicutes;c__Bacilli;o__Lactobacillales;f__La<br>ctobacillaceae;g__Weissella;s__Weissella cibaria | 98.16 | 0.15 | 0 | 97.41 |
| feces_b_bin.14 | d__Bacteria;p__Firmicutes;c__Bacilli;o__Mycoplasmatales;f__<br>_Mycoplasmataceae;g__Spiroplasma_A;s__ | 93.96 | 2.01 | 25 | 83.91 |
| feces_b_bin.12 | d__Bacteria;p__Firmicutes;c__Bacilli;o__Lactobacillales;f__St<br>reptococcaceae;g__Lactococcus;s__Lactococcus lactis_A | 98.3 | 0 | 0 | 98.3 |
| feces_b_bin.10 | d__Bacteria;p__Firmicutes;c__Bacilli;o__Lactobacillales;f__E<br>nterococcaceae;g__Enterococcus_B;s__Enterococcus_B<br>faecalis_A | 94.26 | 0.37 | 0 | 92.41 |
| bran_bin.7 | d__Bacteria;p__Proteobacteria;c__Gammaproteobacteria;o__<br>_Enterobacterales;f__Enterobacteriaceae;g__LSJC7;s__ | 75.68 | 0.58 | 0 | 72.78 |
| bran_bin.3 | d__Bacteria;p__Proteobacteria;c__Gammaproteobacteria;o__<br>_Enterobacterales;f__Enterobacteriaceae;g__Enterobacter;s__<br>_Enterobacter oligotrophicus | 94.86 | 1.48 | 47.37 | 87.46 |
| bran_bin.2 | d__Bacteria;p__Proteobacteria;c__Gammaproteobacteria;o__<br>_Enterobacterales;f__Enterobacteriaceae;g__LSJC7;s__LSJC7<br>sp005671395 | 99.01 | 1.74 | 15 | 90.31 |
| bran_bin.11 | d__Bacteria;p__Firmicutes;c__Bacilli;o__Mycoplasmatales;f__<br>_Mycoplasmataceae;g__Spiroplasma_A;s__ | 74.61 | 2.01 | 0 | 64.56 |
| bran_bin.10 | d__Bacteria;p__Proteobacteria;c__Gammaproteobacteria;o__<br>_Enterobacterales;f__Enterobacteriaceae;g__Mixta;s__Mixta<br>tenebrionis | 99.42 | 2.76 | 18.92 | 85.62 |

Screening MAGs for inferred functions associated with polystyrene and styrene degradation revealed that genes encoding triacylglycerol lipase, an enzyme that could be involved in the breakdown of partially degraded PS (see “*Detection of potential polystyrene and styrene degrading enzymes*”), were only present in MAGs from three classes (Gammaproteobacteria, Actinomycetia, Bacteroidia) recovered from the PS group and the PS feces (**Table S15**). Within the Gammaproteobacteria, all triacylglycerol lipase encoding MAGs were assigned to species in the family Pseudomonadaceae, including *Pseudomonas aeruginosa and Pseudomonas_E qingdaonensis* (**Table S15**). MAGs from these taxa also possessed genes for serine hydrolase (**Table S16**), an enzyme associated with carbonyl group-targeted PS degradation (Kim et al., 2020), for homologues of *styE* (**Table S14**), involved in styrene transport, and for glycol dehydrogenase (**Table S15**), a key enzyme in the direct ring cleavage (**Fig. 5**). Based on these results, we conclude that Pseudomonadaceae strains are promising targets for future studies investigating polystyrene and styrene degrading enzymes and pathways. This conclusion is supported by reports of *P. aeruginosa* strains growing on expanded polystyrene films without additional carbon sources, (Atiq et al., 2010) and of *P. aeruginosa* strains degrading PS-poly lactic acid composites (Shimpi et al., 2012). Furthermore, *P. aeruginosa* strains have been shown to thrive on crude oil (Talaiekhozani et al., 2015) indicating that this species is able to break down a range of hydrocarbons.

Other MAGs possessing lipase genes were assigned to the Bacteroidota genus *Sphingobacterium* and to the Actinobacteria genera *Rhodococcus* and *Corynebacterium*. MAGs from the latter genus also encode *styA* (**Table S15**), the styrene monooxygenase mediating the first step in the vinyl side chain degradation (**Fig. 5**). Several *Corynebacterium* strains, including one identified as *C. pseudodiphtheriticum* based on morphological and physiological properties, have been isolated on styrene, and the authors report that styrene monooxygenase (*styA*) and styrene isomerase (*styC*) are located in the membrane of *Corynebacterium* sp. (Itoh et al., 1996). Therefore, members of this genus should not only be considered as opportunistic pathogens (see “*Microbial community composition*”), but rather be included in future investigations of microbial polystyrene degrading pathways.

### Alternative microbial carbon sources during PS feeding trials

Styrofoam, i.e. the white expanded polystyrene foam (EPA) used as PS source in our study, is manufactured by adding an expanding agent, in most cases the alkane pentane (CCHCC), and the flame retardant hexabromocyclododecane (HBCD) (Law et al., 2005). The EPA used in our study contained 4-7% pentane and up to 1% HBCD at the point of production, according to the manufacturer (see *Material & Methods*). Therefore, we investigated if these compounds could function as carbon or energy sources for the microbial superworm gut community.

The flame retardant HBCD is a persistent organic pollutant that has been detected in a diverse range of environments, wildlife, and humans, and causes developmental neurotoxicity in animals (Szabo, 2014). Several bacterial strains, including *Pseudomonas sp*. HB01, *P. aeruginosa* HS9, and *Rhodopseudomonas palustris* were reported to effectively degrade HBCD (Li et al., 2021). While most chemical compounds produced during bacterial HBCD degradation have been studied in detail (Huang et al., 2019; Wang et al., 2019), only two enzymes, a dehydrogenase and a transferase, involved in catalyzing these reactions have been identified in *R. palustris* (Wang et al., 2019). We did not detect genes encoding 2-haloacid dehydrogenase (K01560) in our data but found genes for glutathione S-transferase (K00799) in all feeding trials and feces samples (**Table S9**). Since both enzymes are thought to work together sequentially (Wang et al., 2019), we conclude that an *R. palustris-*like pathway of HBCD degradation might not be present in the superworm gut microbiome. The presence of glutathione S-transferase genes in all our samples further suggests that this glutathione transferase is widely distributed in prokaryotes and is involved in a variety of processes, as suggested previously (Allocati et al., 2009). However, further transcriptomics and proteomics analysis is required to test for microbial HBCD degradation capabilities and their implications on PS feeding trials of insect larvae.

The ability to degrade alkanes, which include the expanding agent pentane, has been reported for several *Pseudomonas* strains, and enzymes implicated in this process include alkane monooxygenase (*alkB*), cytochrome P450 monooxygenase, and flavin-binding monooxygenase (*almA*) (Liu et al., 2014, p.; Rojo, 2009; Vomberg and Klinner, 2000). Alkane degradation pathways usually start with the oxidation of a terminal methyl group yielding a primary alcohol, which is further oxidized to an aldehyde, converted into a fatty acid, and finally oxidized through beta-oxidation to render CO_2_ (Rojo, 2009). We did not detect any of the involved monooxygenase genes in our samples, except for *alkB* genes in the PS group and the PS feces (**Table S17**). The absence of a complete pentane degradation pathway in the superworm gut could be due to the limited oxygen availability in the intestine, since all aforementioned alkane degradation pathways rely on oxygen for substrate activation and as the terminal electron acceptor (Rojo, 2009). Anaerobic microbial alkane degradation pathways exist and have been reported to involve novel hydrocarbon activation mechanisms (Widdel and Rabus, 2001), although the involved enzymes remained unexplored. To our knowledge, the only anaerobic activation of alkanes studied on a transcriptional level involved *Desulfatibacillum alkenivorans* AK-01 (Herath et al., 2016). The proposed key enzyme is an alkylsuccinate synthase (*ass*), which initiates the alkane degradation via the addition of a fumarate molecule. *Ass* homologues were detected in the bran, PS, and starvation feeding trials, and in the bran and PS fecal samples (**Table S17**). Based on our findings, it is possible that members of the superworm gut microbiome use pentane as a carbon source or for energy conservation. However, pentane availability in styrofoam is likely limited. Pentane diffuses rapidly after manufacture to less than 1% (wt) content over a period of about 2–3 weeks, and the remaining pentane continues to diffuse to insignificant levels within several weeks after production (Simpson et al., 2020). Regardless, further studies are needed to explore the abilities of microbial growth on alkenes, such as pentane, in the superworm gut.

## Conclusion and Outlook

Our study provides the first metagenomic analysis of the gut microbiome from the superworm (*Zophobas morio*), comparing microbial communities of worms reared on PS, on regular bran feed, and under starvation conditions. The diet consisting solely of PS had considerable influences on the microbiome beyond the detection of PS degradation capabilities. It was characterized by a loss of microbial diversity, the detection of opportunistic pathogens indicative of dysbiosis, and an enrichment of inferred functions for stress response. These modifications of the gut microbiome, in combination with the minimal weight gain of the superworms reared on PS, paints a picture of survival under poor health conditions for the insect host. The presence of inferred polystyrene and styrene degrading functions, supports previous studies reporting polystyrene degrading abilities of the superworm microbiome. However, we cannot rule out that the microbes also metabolized other styrofoam components, such as the flame retardant or the remains of the expanding agent.

Our results support previous suggestions that superworms can help to reduce PS waste (Kim et al., 2020). However, the minimal weight gain of the larvae on a PS diet will likely hamper their use in the polystyrene recycling process. In particular, downstream applications such as biodiesel production from superworm fatty esters, an approach that has been proven feasible using superworms raised on regular feed (Leung et al., 2012), might not be achievable. Diet diversification, for example by supplementing styrofoam with food waste, could help to counteract the dietary deficits of the unbalanced PS feed and might increase gut microbiome health and subsequently host weight gain. Such an approach could also contribute towards waste valorization of agricultural and industrial by-products by insects (Derler et al., 2021). Alternatively, isolating PS degrading microbes, and characterizing their enzymes involved in PS degradation pathways, followed by enzyme engineering and large-scale production, are possible options to utilize the superworm microbiome.

In summary, our metagenomic exploration of the superworm gut microbiome provided the first insights into how their microbial community responds to a PS diet. However, comparable to opening a can of worms, our results also raised many new questions. Which members of the microbial community are active, and which genes are transcribed during a PS diet compared to regular feed? What are the complete pathways of polystyrene and styrene degradation used by the gut microbes? Can some bacteria conserve energy by using styrofoam components other than polystyrene? Employing transcriptomics to quantify gene expression levels (Hawley et al., 2017), and using click chemistry to visualize *in situ* changes of translational activities (Lindivat et al., 2020) will contribute to answering these questions. However, bringing the majority of superworm gut bacteria into culture remains the ultimate goal in our endeavor to characterize microbial PS degradation in this and other insect microbiota.

## Abbreviations

PS: polystyrene
EPA: expanded polystyrene foam
HBCD: Hexabromocyclododecane.

## DATA SUMMARY

All sequencing data has been deposited with the National Center for Biotechnology Information (NCBI) under BioProject accession number **PRJNA801070**. Supplementary material can be found on Figshare: https://figshare.com/xxxxx.

## Acknowledgements

We thank Phil Hugenholtz and Gene Tyson for their help in obtaining financial support for the project leading to this publication, and the ACE team (http://ecogenomic.org) for stimulating discussions.

## Authorship and contributions

Conceptualization by C.R.; Investigation by J.S. and C.R.; Formal Analysis by J.S., A.P., and C.R.; Data Curation by J.S.; Writing – Original Draft Preparation by J.S. and A.P.; Writing – Review and Editing by S.A. and C.R.; Visualization by J.S., A.P., and C.R.; Supervision by C.R.

## Suppl. Figures

**Figure S1.**
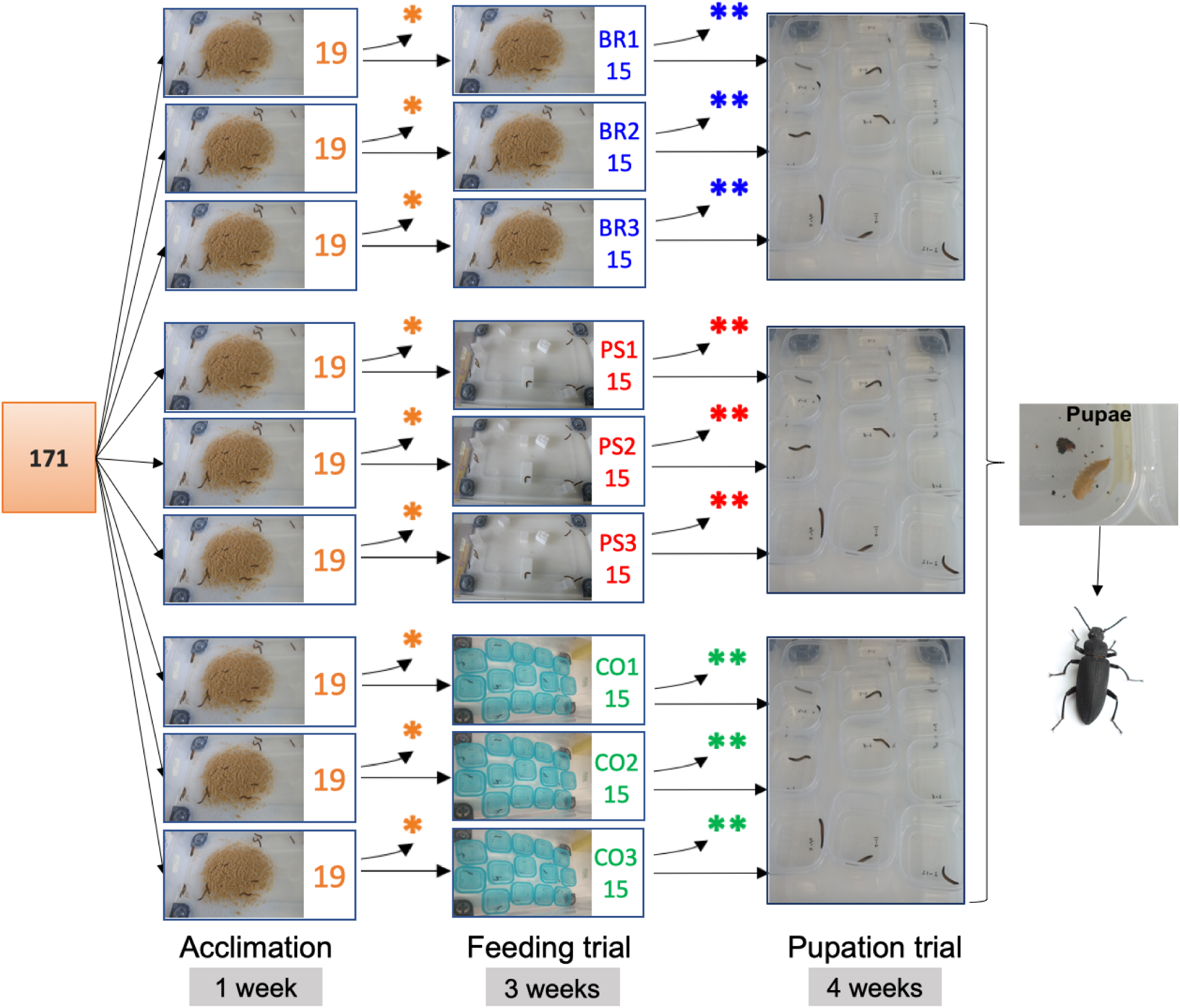
Experimental layout of the superworm polystyrene diet experiment. In total, 171 worms were kept in nine containers during the acclimatization period of one week and fed with bran and carrots. At the start of the feeding trial the larvae were divided into three groups, each with three replicates (containers), which were labelled according to the food source: i) The bran group (**BR1, 2, 3**) was feed solely with bran, ii) the polystyrene group (**PS1, 2, 3**) had solely PS as food source and iii) the control group (**CO1, 2, 3**) received no food supply. Feeding for the BR and PS group occurred once a week. All three treatments were accompanied by misting the lid of the container with water to prevent the worms from dehydrating. Asterisks indicate the processing of larvae for microbiome analysis: * post acclimation period, 4 worms were randomly selected and removed from each container, one worm from each container was randomly selected for microbiome sequencing; ** post-feeding trial analysis, 1 worm from each container was randomly selected for microbiome sequencing.

**Figure S2.**
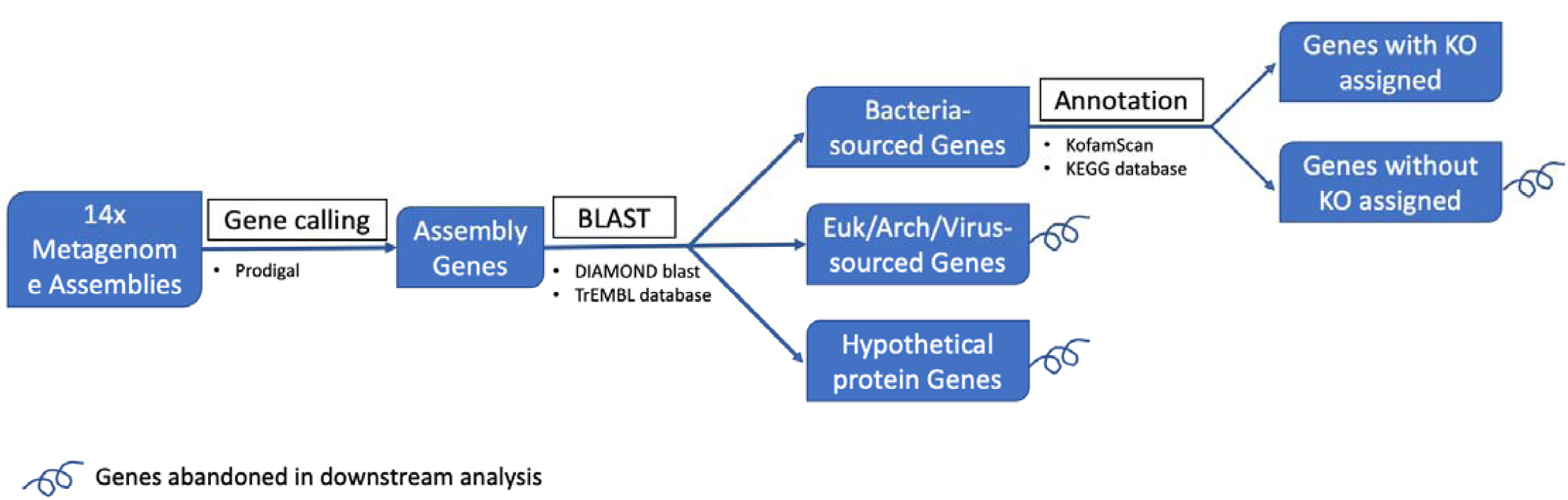
Annotation of bacterial genes. Genes with a bacterial top blast hit were selected and annotated by KOfamScan for gene profiling analysis, while eukaryotic/ archaeal/ viral genes and genes without blast hits were excluded from the downstream analysis. “Hypothetical protein genes” are genes that did not have a blast hit.

**Figure S3.**
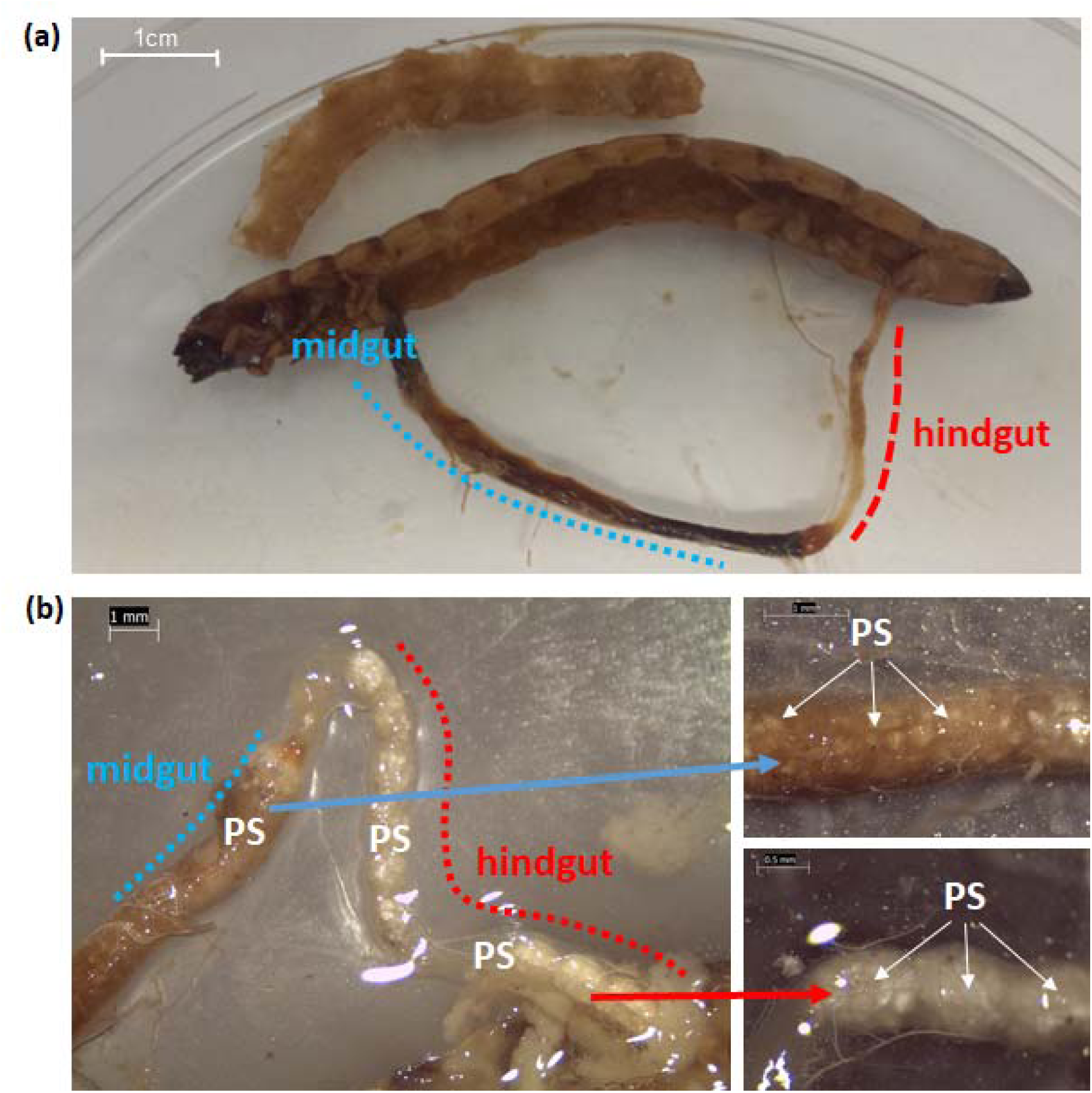
Superworm anatomy and gut content. (a) Dissection of a superworm (*Zophobas morio*). Midgut and hindgut are highlighted. The removed ventral section is shown above the larvae. (b) Sections of mid and hindgut tightly packed with polystyrene (PS) particles. Arrows pointing to higher magnification images of the PS packed midgut (upper image) and hindgut (lower image).

**Figure S4.**
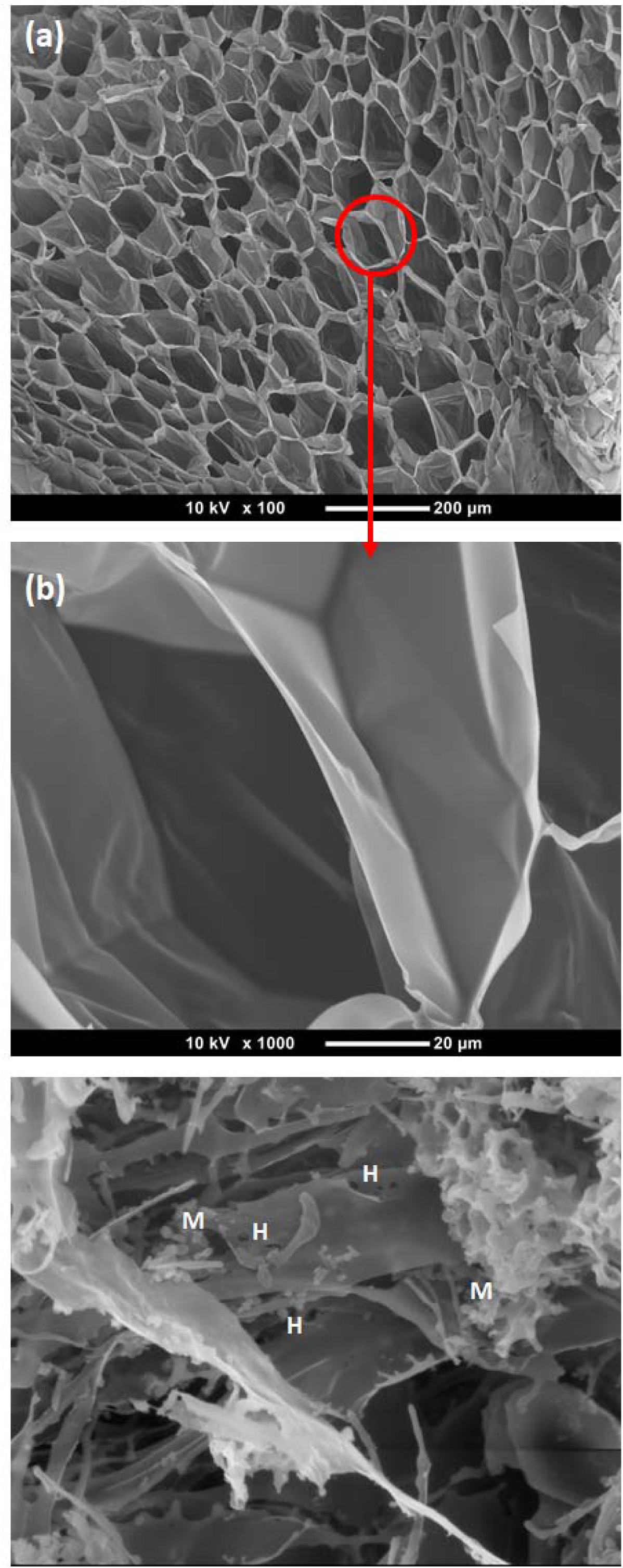
Scanning electron pictures of polystyrene particles. (**a**) Scanning electron microscopy (SEM) image of the extruded polystyrene (EPS; styrofoam) as supplied by the manufacturer. (**b**) A higher magnification of the same particle (position in (a) is indicated by a red circle) showing the clean and intact PS foam structure. (**c**) Polystyrene particle recovered from the gut of a superworm in the PS group. Note the rugged structure of the PS, several holes (H), and microbial cells (M).

**Figure S5.**
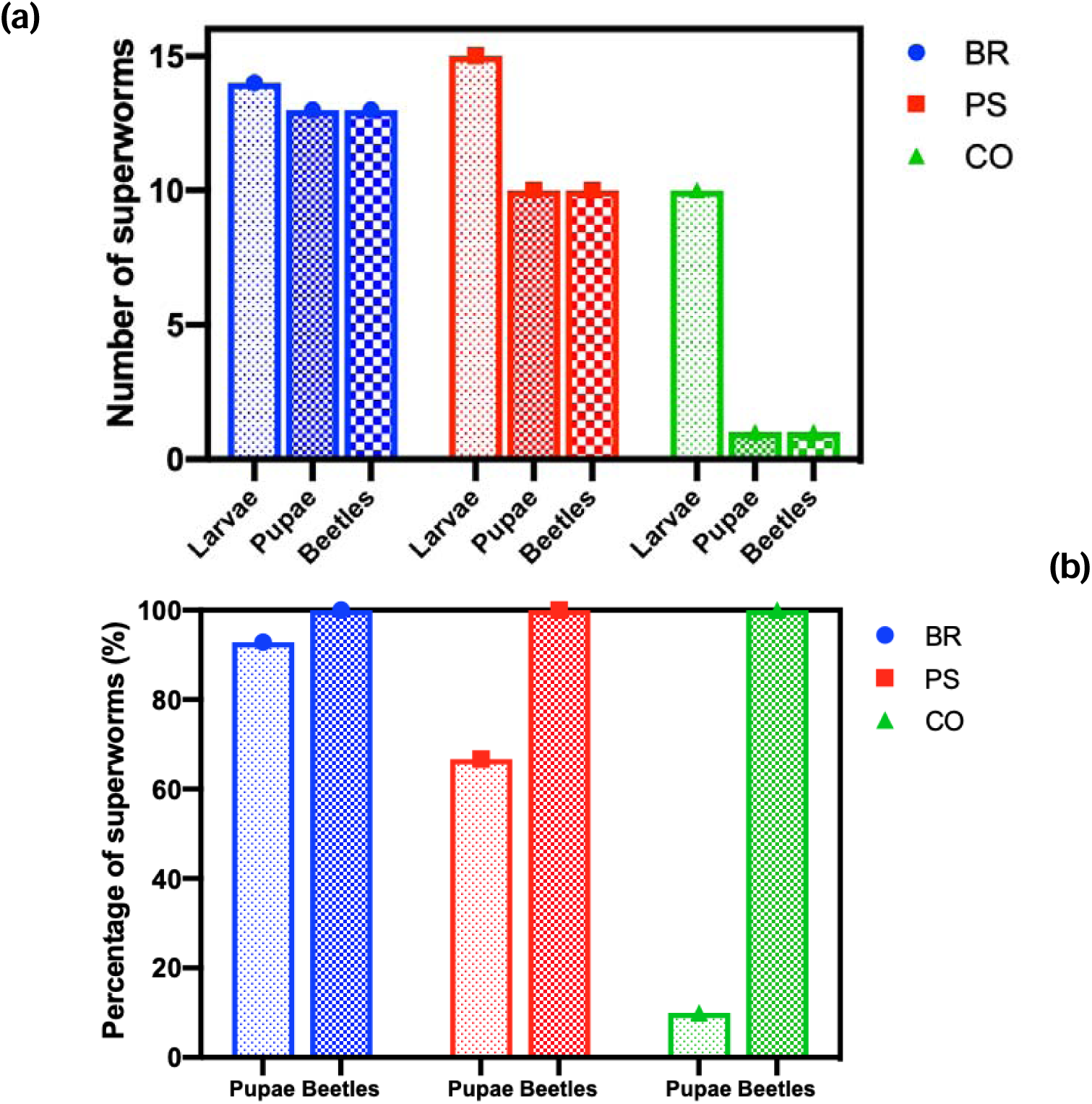
Superworm pupation experiment. Numbers **(a)** and percentages **(b)** of superworms (Larvae) entering the pupae phase (Pupae) and emerging as live imago (Beetles) post transformation, are provided for all three feeding trials. Note that in (b) the percentage of pupae has been calculated by setting the number of larvae in each experiment to 100%, and the percentage of beetles has been calculated by setting the number of pupae in each experiment to 100%. Abbreviations: BR = bran group; PS = polystyrene group; CO = starvation control group.

**Figure S6.**
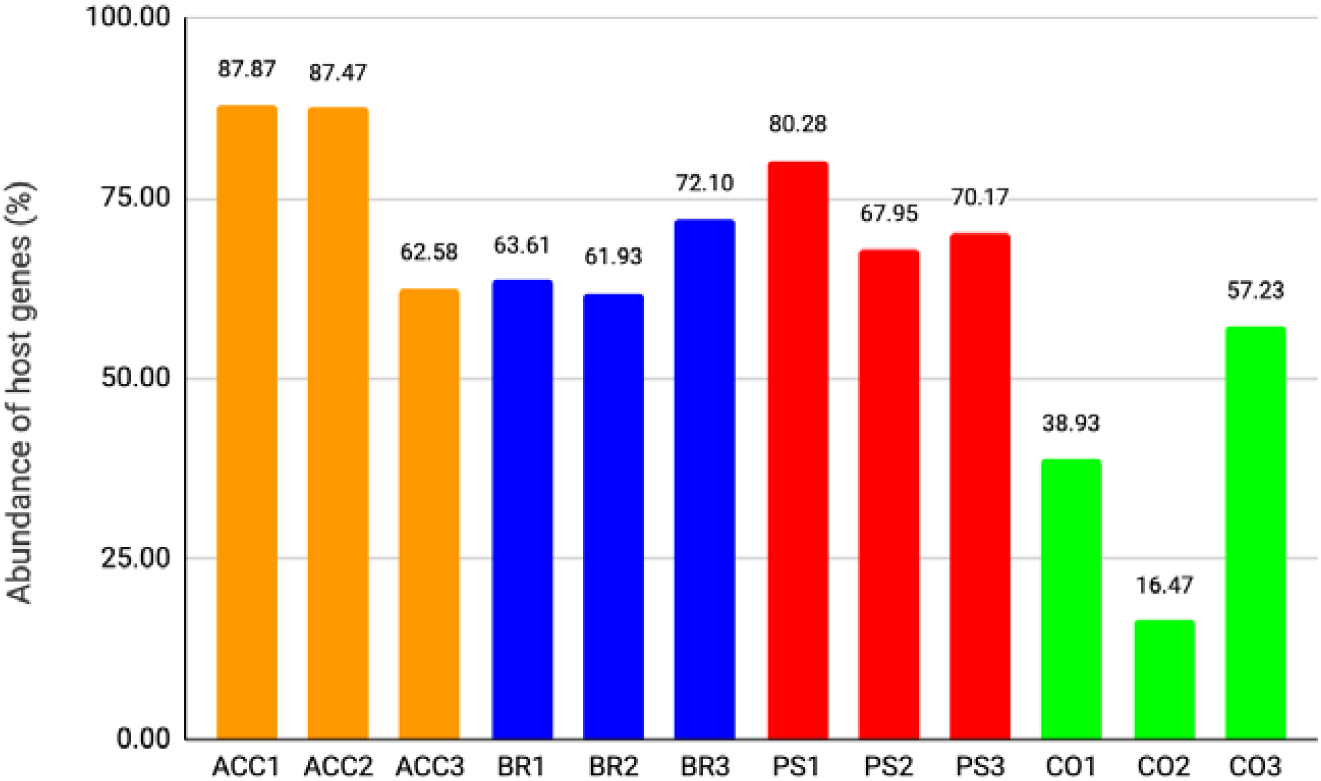
Estimated proportion of superworm host genes in metagenomes. Host gene proportions were calculated by dividing the number of 18S rRNA reads assigned to ‘p Arthropoda’ by the number of total rRNA (16S and 18S) reads based on read assignments by graftM using the custom package “4.40.2013_08_greengenes_97_OTUs_with_euks.gpkg”. Abbreviations: ACC: Acclimation period (pre feeding trials); BR: Bran group; PS: PS group; CO: control starvation group.

**Figure S7.**
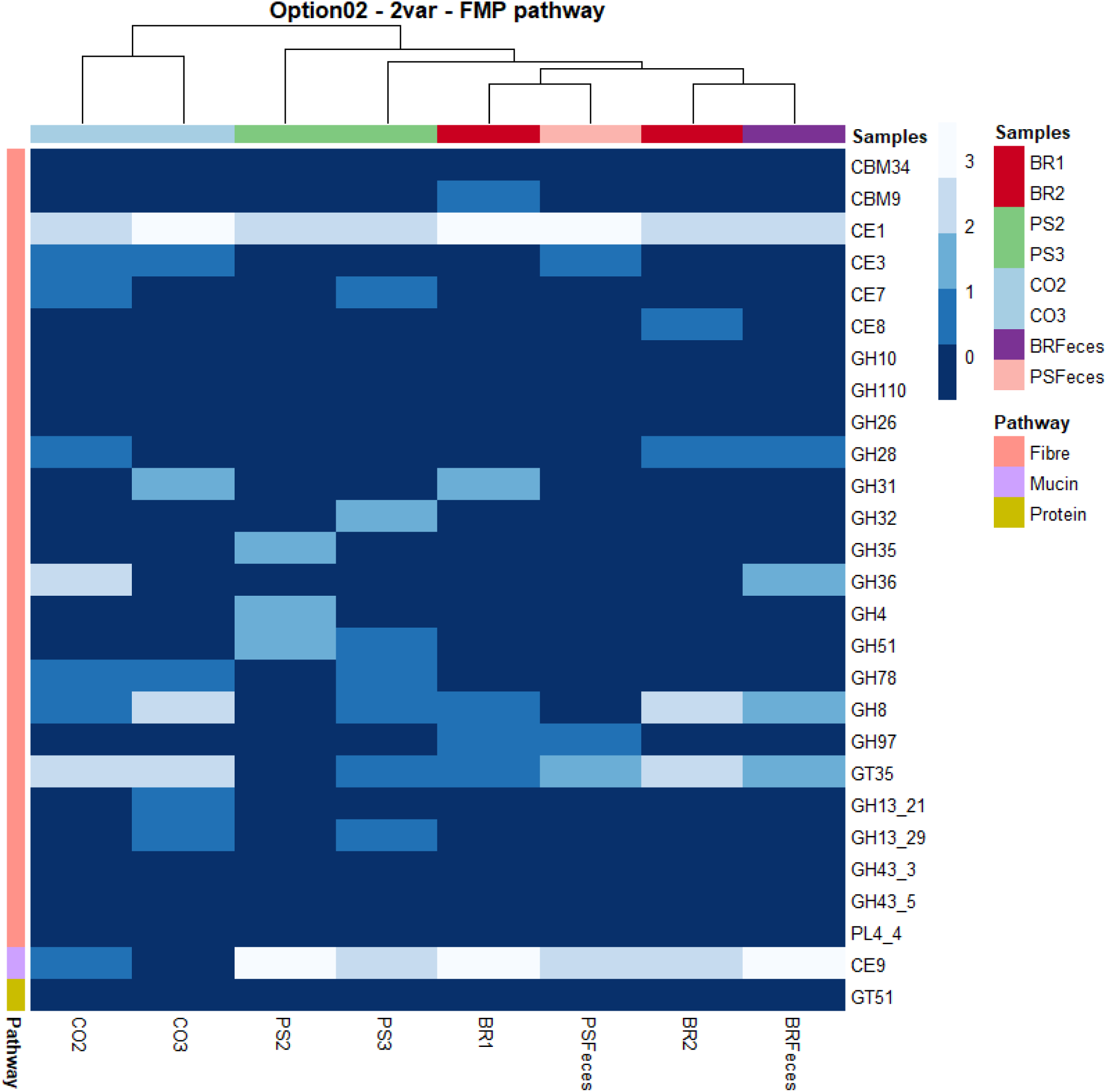
Bacterial genes involved in fiber, mucus, and protein degradation, annotated against CAZy. The heatmap includes samples from the bran group (BR), the polystyrene group (PS), the starvation control group (CO), the polystyrene feces (PSF), and the bran feces (BRF). The y-axis is sorted by fiber, mucus, and protein degrading abilities.

**Figure S8.**
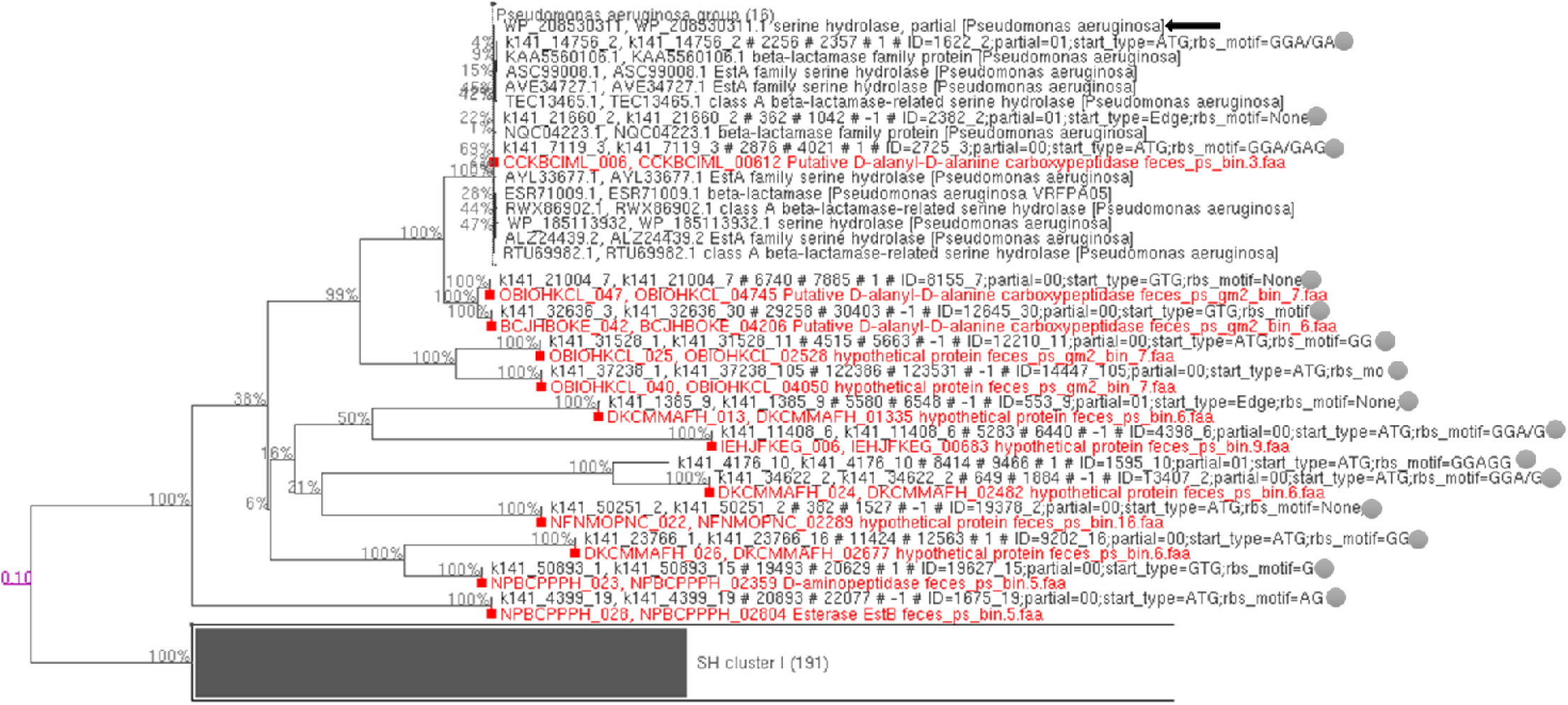
Phylogenetic tree of serine hydrolase genes. Homologues of WP_208530311.1, the gene encoding serine hydrolase in *Pseudomonas aeruginosa* (black arrow), were retrieved from assemblies and MAGs by a blastp search with evalue <1e-5. The phylogenetic tree was inferred with IQ-TREE (LG+C10+F+G+PMSF model) from a 998-position alignment with true bootstrap support values (gray numbers at internal nodes) based on 100 trees inferred under the same model. The tree was rooted between the group containing the reference serine hydrolase sequence and the rest. Fifteen sequences from assemblies (gray dots) and twelve from MAGs (red font) of sample Feces-PS and PS1 were clustered with WP_208530311.1, suggesting they represent true homologues of serine hydrolase genes and could translate into a protein involved in the breakdown of polystyrene.

**Figure S9.**
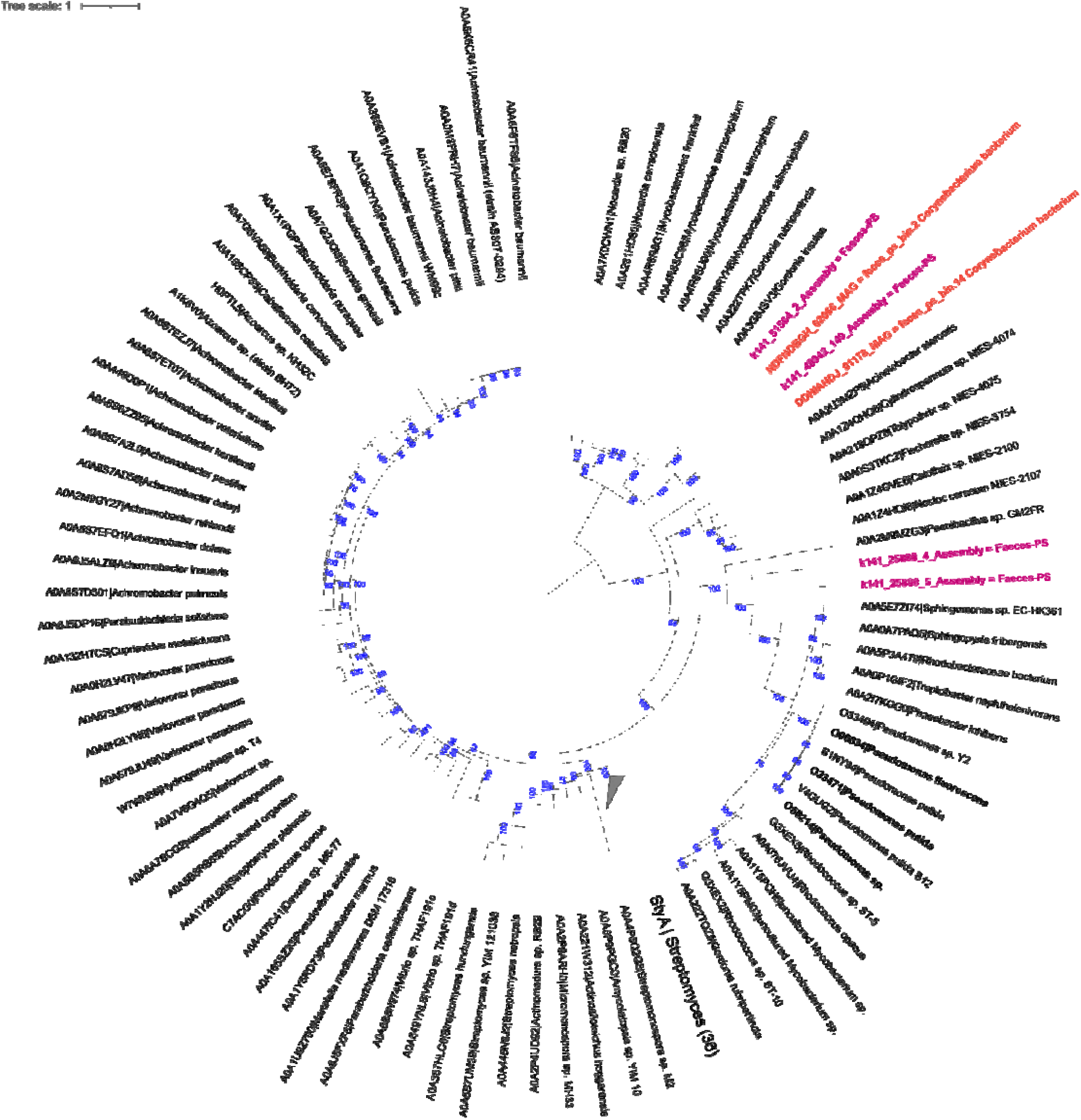
Phylogenetic tree of styA gene homologues encoding styrene monooxygenase. The phylogenetic tree was inferred with IQ-TREE (LG+C10+F+G+PMSF model) from a 787-position alignment with ultrafast bootstrap support values (blue numbers at internal nodes) based on 1000 ultrafast trees under the same model. The tree was rooted on the group containing two StyA sequences from MAGs (red) and two StyA from Faeces-PS assembly (pink). The styA genes of the experimentally verified enzymes in *Pseudomonas fluorescens, Pseudomonas putida*, and *Pseudomonas sp*. are highlighted in bold.

**Figure S10.**
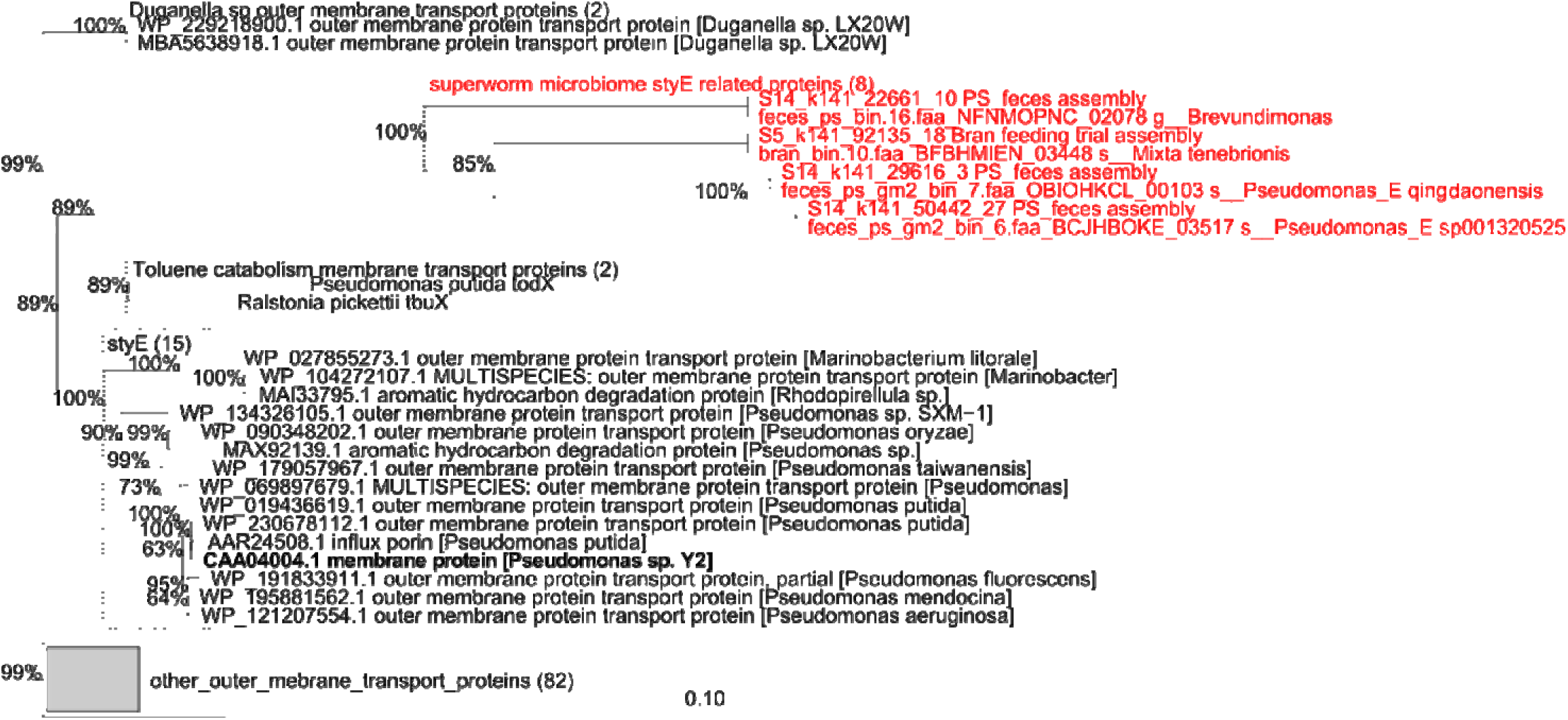
Phylogenetic tree of styE gene homologues encoding an outer membrane protein involved in styrene transport. Homologues of the StyE encoding gene from Pseudomonas sp. Y2 (highlighted in bold) were retrieved from NCBI (blastp; 100 best hits), and were recovered from the superworm microbiome assemblies and MAGs (blastp, <e-10; highlighted in red). In addition, two genes, which represent membrane proteins involved in toluene catabolism, from the INTERPRO family “Outer membrane protein transport protein (OMPP1/FadL/TodX) IPR005017”, which includes styE, were added. The phylogenetic tree was inferred with IQ-TREE (LG+C10+F+G+PMSF model) with ultrafast bootstrap support values based on 100 ultrafast trees inferred under the same model.

**Fig. S11.**
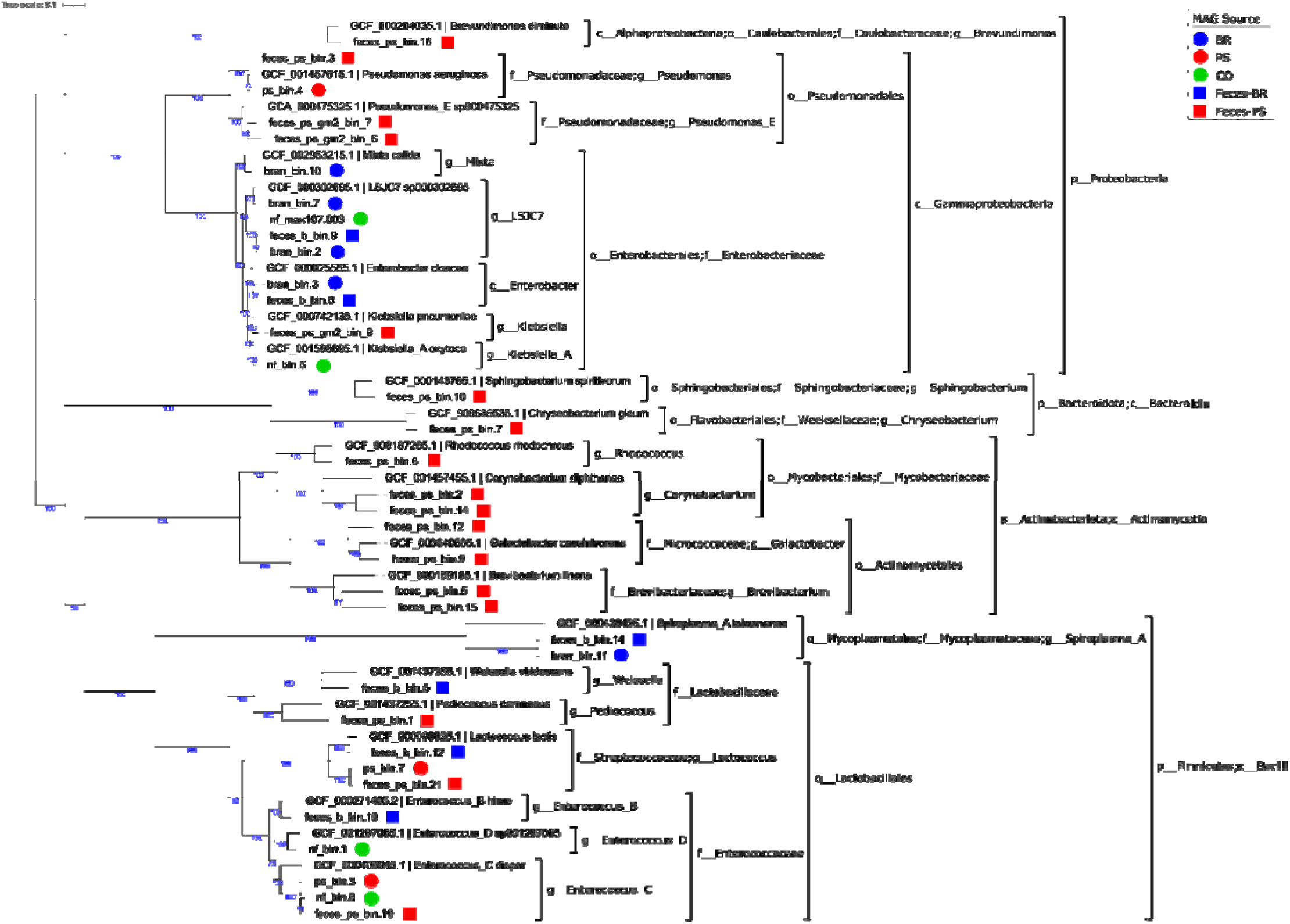
Phylogenetic tree of metagenome-assembled genomes (MAGs) recovered in this study. A multiple sequence alignment of 56 genomes (35 MAGs plus 21 GTDB species-representative genomes selected from the same genera as the MAGs) was created based on 120 marker genes using gtdb-tk. Starting trees were inferred with FastTree under the JTT+CAT model and used for IQ-TREE under the LG+C10+F+G+PMSF model. 100 bootstrap trees were calculated under the same IQ-TREE model, and bootstrap values above 70% are shown on the corresponding branches.

